# De novo protein design in an expanded chemical space

**DOI:** 10.64898/2026.10.08.757794

**Authors:** Yuxuan Li, Yanbo Jing, Yeyu Su, Wei Ren, JiaYi Zhang, Tao Liu

**Affiliations:** State Key Laboratory of Natural and Biomimetic Drugs, Chemical Biology Center, Department of Molecular and Cellular Pharmacology, School of Pharmaceutical Sciences, Peking University, 38 Xueyuan Road, Haidian District, Beijing 100191, China

## Abstract

Generative protein design has made sequence and structure programmable, yet its chemical alphabet remains largely confined to the canonical twenty. Here, we establish CheMoDesign, an experimentally validated generative framework for designing proteins *de novo* around user-specified chemistry. CheMo Design weakens pair-representation conditioning in a pretrained all-atom structure-prediction model while preserving chemical anchors, then refines generated backbones through sequence-conditioned denoising. By treating post-translational modifications, synthetic residues, and chemical modifications as user-defined building blocks, the framework generated proteins with chemistry-dependent recognition and reactivity in single-round experimental campaigns. Sulfotyrosine and nitrotyrosine enabled modification-dependent recognition of thrombin and HER2, respectively. Genetically encoded noncanonical residues enabled bioorthogonal affinity switching and light- or proximity-triggered covalent target capture. Post-expression chemical modifications yielded a light-activated covalent PD-L1 binder and an isoform-selective CA IX inhibitor (K_D_ = 45 nM) designed to engage catalytic Zn²⁺. CheMoDesign makes chemical mechanism a programmable dimension of *de novo* protein design.

## Main Text

Generative models can now produce novel protein sequences and structures and increasingly support the design of functional proteins and protein–protein interactions (*1–5*). Yet contemporary de novo protein-design workflows remain built predominantly around the canonical amino-acid alphabet (*6, 7*). Their functional repertoire is therefore constrained by the chemistry that canonical side chains can execute. Natural proteins partly escape this limit through post-translational modifications, cofactors, and chromophores that enable regulated conformational changes, catalysis, and signal conversion (*8, 9*). Synthetic chemistry offers an even broader range of reactions and physical properties that biosynthesis and natural evolution have not explored. These chemical capabilities are not routinely accessible as explicit variables in generative protein design, sharply limiting the mechanisms that can be programmed into designed proteins.

Noncanonical amino acids and chemical modifications have long expanded the capabilities of existing proteins (*10–12*). Most current strategies, however, either install noncanonical residues or reactive chemical functionalities into existing protein or peptide frameworks through substitution, conjugation, or local redesign (*7, 13–15*), develop chemistry-specific design protocols around particular functional groups (*16, 17*), generate and experimentally validate ncAA-containing peptides (*18–20*), or design proteins around separate small-molecule ligands (*21*). These studies show that noncanonical chemistry can be accommodated in designed molecules, but do not yet provide a general generative framework that actively exploits its chemical properties to build de novo interfaces and functions across diverse targets. The inverse problem is therefore to begin with a desired chemical capability and generate a protein that positions and exploits that chemistry as an integral part of molecular recognition or function. Solving this problem would bring the broader functional repertoire of synthetic chemistry into generative protein design, allowing chemical functionality itself to be specified, exchanged, and programmed as part of the design.

We developed CheMoDesign to address this inverse problem. Without retraining or fine-tuning, the method repurposes the structural prior of the pretrained Boltz-2 all-atom structure-prediction model by selectively weakening its pair-representation conditioning, while preserving user-defined functional anchors and optimizing internal representations with geometric losses. A subsequent sequence-conditioned refinement step reconciles generated backbones with realizable sequences. Atom-level descriptions allow post-translational modifications (PTMs), genetically encoded noncanonical amino acids (ncAAs), and post-expression chemical modifications to serve as organizing elements of the generated interface. We tested the framework across recognition, affinity switching, photoactivation, proximity-driven covalency, and metal-coordination-mediated inhibition of a metalloenzyme. Each experimental campaign comprised a single computational design round without experimental optimization.

### Weakening structural conditioning enables generative protein design

In an AlphaFold 3-like predictor, single and pair representations produced by the Pairformer condition the diffusion module toward the structure implied by the input sequence (*22, 24*). Pair-representation scaling was recently introduced as an inference-time strategy for modulating conformational sampling in AlphaFold 3 and Boltz-2 (*25*). We asked whether controlled weakening of this structural conditioning could instead be repurposed for de novo protein generation while preserving user-defined functional constraints. CheMoDesign scales the pair representation as z_ij_ → (1 − α)z_ij_ while exempting interactions that define a functional anchor (Fig. 1A). Geometric losses are backpropagated through a single denoising step to optimize the modulated single and pair representations. Diffusion from these weakened but functionally constrained representations produces diverse target-associated backbones, a process we term “dreaming.” The same framework accepts atom-level descriptions of PTMs, ncAAs, and post-expression modifications, allowing specified chemical groups to participate directly in generation, computational filtering, and experimental realization (Fig. 1B).

**Fig. 1.**
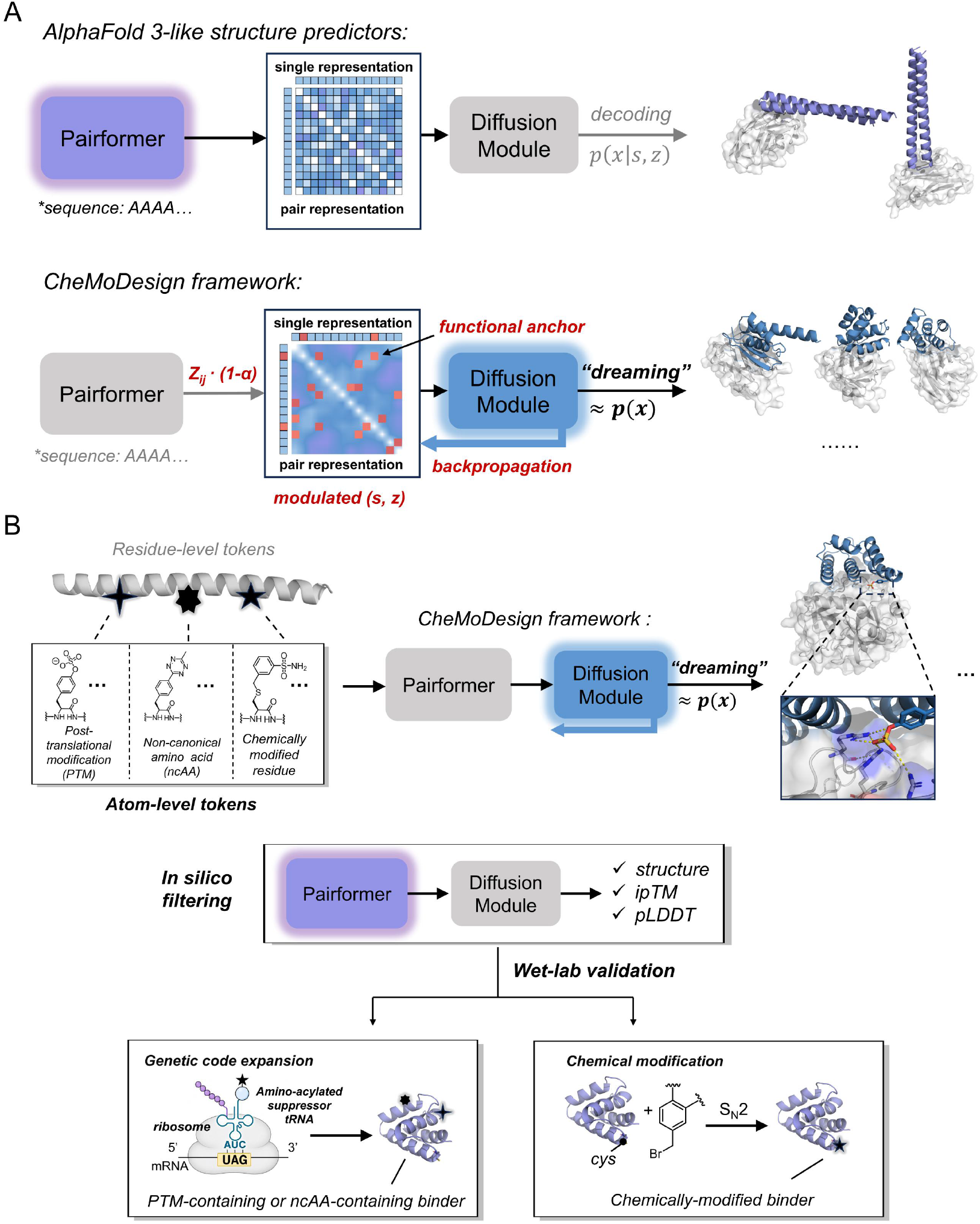
CheMoDesign repurposes structure-prediction models for protein design across expanded chemical space. (A) Conventional AlphaFold 3-like inference decodes structures conditioned on Pairformer-derived single and pair representations. CheMoDesign selectively weakens pair conditioning while preserving functional anchors and optimizes internal representations through geometric-loss backpropagation across one denoising step. Diffusion from the modulated representations samples diverse candidate structures (“dreaming”). (B) Atom-level representations of PTMs, ncAAs, and chemically modified residues are incorporated as specified chemical entities. Designs are filtered computationally and realized by genetic code expansion or post-expression modification.

We implemented CheMoDesign in Boltz-2 (*22*) and first examined pair-representation scaling using PD-L1 as a test target and an 80-residue polyalanine input (Fig. 2A). Among 100 generated backbones at each scaling factor, the number passing compactness and secondary-structure filters increased to 40 at α = 0.9. Four ProteinMPNN (*23*) sequences were then generated per backbone and evaluated by AlphaFold 3 structure prediction. The mean ipTM of the top 10 designs rose markedly with scaling, indicating improved overall interface designability, whereas the number of designs crossing the hard thresholds of minPAE ≤1.5 Å, ipTM ≥0.8, and pLDDT ≥80 remained low and did not increase consistently. Thus, weakening the pair representation released the diffusion prior and improved the quality of the best samples, but did not by itself ensure compatibility between a generated backbone and a discrete amino acid sequence.

**Fig. 2.**
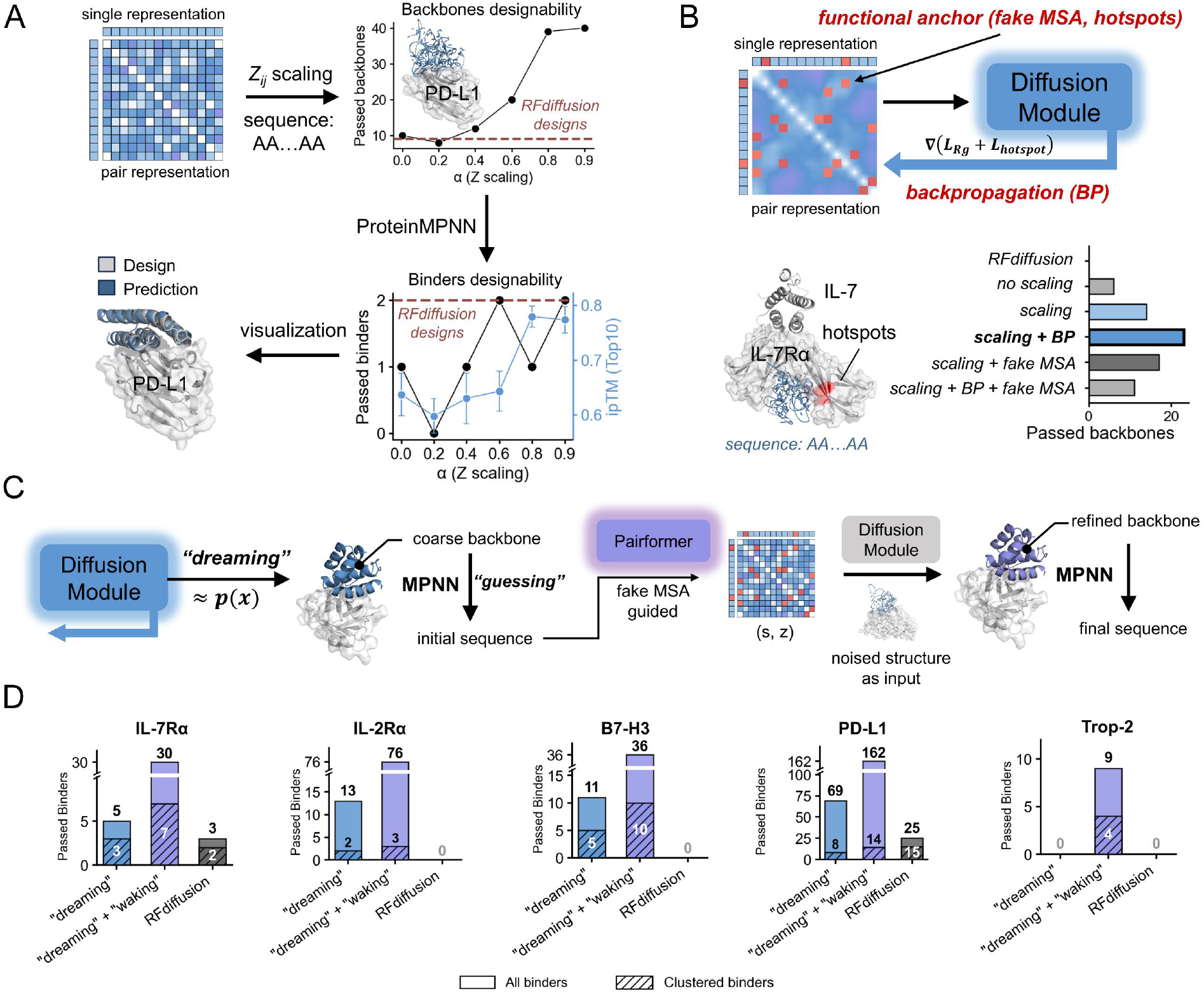
Establishment of the CheMoDesign protein-design workflow. (A) Effect of pair scaling, zij → (1 − α)zij, on PD-L1 backbone and sequence design in Boltz-2. For each α, 100 backbones were generated from an 80-residue polyalanine input. Passing backbones had <40% loop content and radius of gyration <14 Å. Four ProteinMPNN sequences per backbone were evaluated by AlphaFold 3; passing binders had minPAE ≤1.5 Å, ipTM ≥0.8, and pLDDT ≥80. Blue, mean ± SD ipTM of the top 10 designs. (B) Hotspot-directed IL-7Rα design using backpropagation (BP), artificial MSA-derived pair signals (fake MSA), or both. Passing backbones additionally placed at least two binder residues within 7 Å (Cα-Cα) of a specified hotspot. (C) Waking refinement combines an initial MPNN-designed sequence, binder-directed fake-MSA and template features, and a partially noised dreamed structure for truncated denoising and final sequence design. (D) Passing and structurally distinct binders generated by dreaming, dreaming plus waking, or RFdiffusion across five targets. Each condition started from 100 backbones. Passing sequence–backbone pairs were counted without collapsing sequences from the same backbone. Sequence sampling and the RFdiffusion PD-L1 hotspot-filter exception are described in Methods; clusters used a TM-score cutoff of 0.6.

We next asked whether local interface requirements could be retained while global structural conditioning was relaxed. Using IL-7Rα as an example, we designated target hotspot residues as functional anchors whose associated representations were protected from scaling, and introduced additional hotspot guidance through geometric-loss backpropagation (BP), artificial MSA-derived pair features (fake MSA), or both (Fig. 2B and fig. S1). Scaling combined with BP produced 23 of 100 backbones that passed the global compactness and secondary-structure filters and also placed at least two binder residues within 7 Å (Cα-Cα) of the specified hotspot, compared with 14 for scaling alone. RFdiffusion (*2*) produced no passing backbone under the same hotspot and filtering criteria. These results show that local interaction requirements can remain enforceable even when global pair conditioning is weakened.

Strong perturbation of the internal single and pair representations increased structural exploration but could also produce locally inconsistent geometries, including implausible covalent-bond arrangements. We therefore separated the workflow into “dreaming” and “waking” steps (Fig. 2C and fig. S2). After dreaming and initial MPNN sequence design, the sequence, binder-directed fake-MSA and template features, and a partially noised dreamed backbone were supplied to truncated diffusion denoising. This sequence-conditioned waking step repaired local geometry and yielded a refined backbone for final sequence design. Across IL-7Rα, IL-2Rα, B7-H3, PD-L1, and Trop-2, waking increased the number of computationally passing binders from 0 to 9, with the sharpest gain for Trop-2 (Fig. 2D). Each target retained 3 to 14 structurally distinct clusters at a TM-score cutoff of 0.6, whereas RFdiffusion produced 0 to 25 passing binders per target from 100 initial backbones per target, with branch-specific sequence-sampling budgets described in Methods. Thus, dreaming supplies structural exploration, and waking converts that exploration into sequence-compatible, locally coherent designs.

### Chemical anchors organize generated protein interfaces

We reasoned that an ncAA-containing protein can be viewed simultaneously as a polymer and as a protein–ligand complex: the canonical backbone follows protein-level priors, whereas the noncanonical group can draw on chemical-recognition priors learned from small molecules. We therefore retained the residue-level representation of the parent amino acid and appended an atom-level representation of the noncanonical chemical group, linked according to the corresponding covalent-bond topology (Fig. 3A). Every pair involving the atom-level chemical component was treated as an anchor pair and protected during scaling and gradient optimization, preserving its relationship with the protein context. LigandMPNN (*26*) was used for sequence design and waking refinement. This construction allowed the same framework to operate on PTMs, ncAAs, and residues installed by post-expression chemistry.

**Fig. 3.**
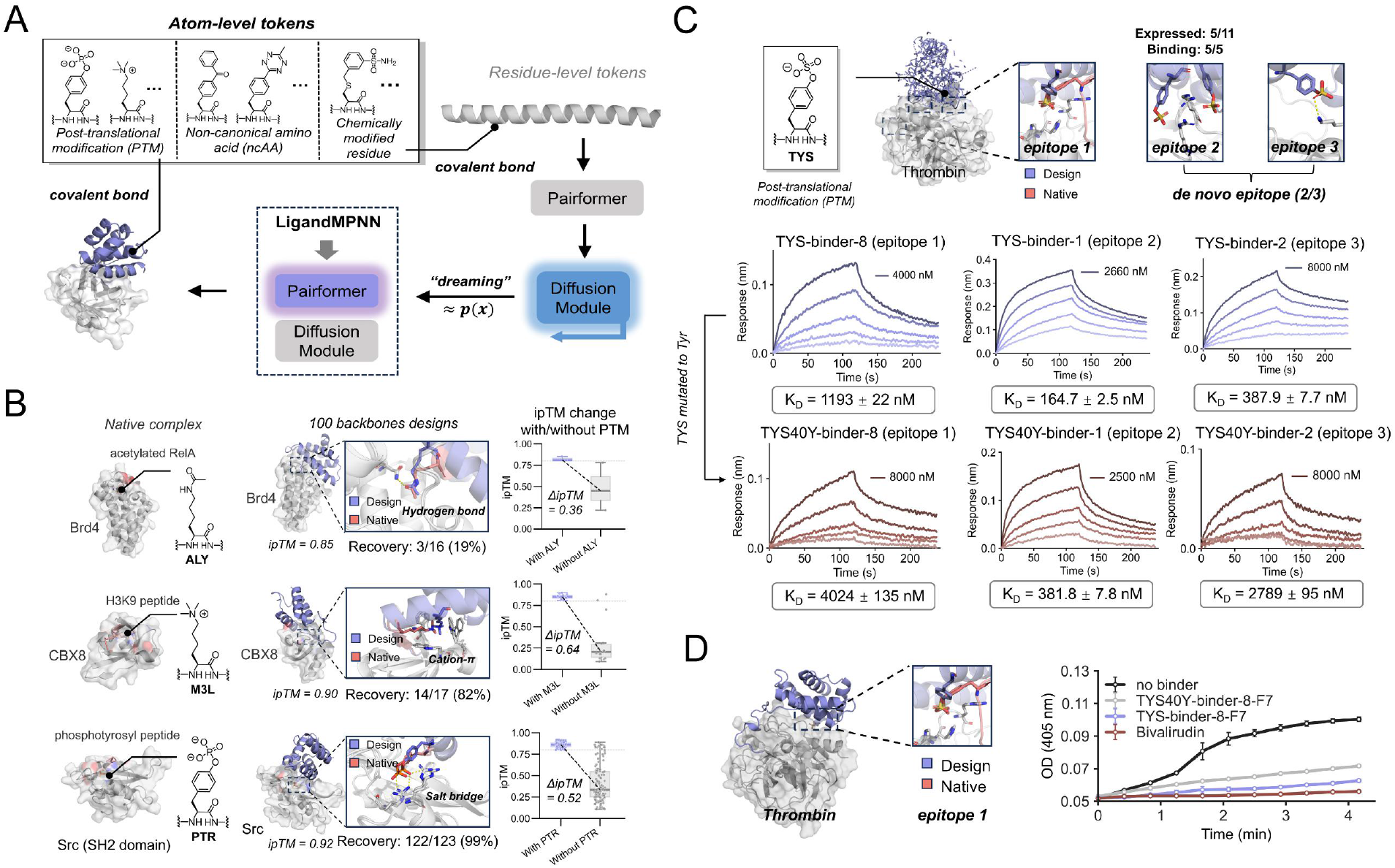
CheMoDesign enables protein design with noncanonical chemical functionalities. (A) Residue-level and atom-level representations of PTMs, ncAAs, and chemically modified residues are linked; pairs involving the atom-level chemical component are preserved as anchors during scaling and gradient optimization. LigandMPNN is used for sequence design and waking. (B) Recovery of PTM-dependent interactions in native complexes containing acetyllysine (ALY), trimethyllysine (M3L), or phosphotyrosine (PTR). Recovery required a passing design and PTM functional-group RMSD <2 Å from native geometry. Chemical-ablation controls were converted to the corresponding canonical residue and repredicted. (C) TYS-containing thrombin binders at three epitopes and matched TYS40Y variants characterized by BLI. Five of 11 selected designs expressed; all five bound thrombin. KD values and uncertainties are from 1:1 kinetic fits. (D) Raw OD405 trajectories from a chromogenic thrombin assay for TYS-binder-8 fused to a C-terminal TTI-derived active-site-targeting fragment. Constructs and bivalirudin were tested at 5 μM. Data are mean ± SD from three technical replicates.

As a retrospective test, we asked whether CheMoDesign could recover modification-dependent interactions from native complexes containing acetyllysine, trimethyllysine, or phosphotyrosine (Fig. 3B and fig. S3). Among 100 generated backbones per complex, native-like chemical geometry was recovered in 3 of 16, 14 of 17, and 122 of 123 computationally passing designs, respectively, using a functional-group RMSD threshold of 2 Å. Converting each modification to its canonical parent residue and repredicting the same design reduced ipTM by 0.36, 0.64, and 0.52, respectively. The unequal effects were consistent with different degrees of chemical dependence across the three native interfaces, while demonstrating that the framework could preserve interactions arising specifically from the modified group.

We then prospectively designed sulfotyrosine (TYS)-containing binders to thrombin. Although generation was initialized with specified hotspot residues, we did not discard backbones that migrated to other regions of the target. CheMoDesign consequently recovered a known TYS-compatible region and, to our knowledge, predicted two additional TYS-compatible binding modes at regions not represented by known TYS complexes (Fig. 3C; 27, 28). Five of 11 selected designs were expressed, and all five bound thrombin. Representative designs from the three epitopes bound with dissociation constants of 1193 ± 22 nM, 165 ± 3 nM, and 388 ± 8 nM. Replacing TYS40 with Tyr weakened binding to 4024 ± 135 nM, 382 ± 8 nM, and 2789 ± 95 nM, respectively. The magnitude of dependence varied by interface, but all three matched comparisons favored the designed modified residue.

To connect chemical recognition to function, we fused TYS-binder-8 to a C-terminal active-site-targeting fragment derived from the tsetse thrombin inhibitor (TTI) (Fig. 3D). At 5 μM, TYS-binder-8-F7 suppressed chromogenic substrate turnover more strongly than the matched TYS40Y-binder-8-F7 variant, whereas the no-binder control showed an unimpeded increase in OD405 and 5 μM bivalirudin nearly abolished activity (mean ± SD, n = 3 technical replicates). Because the reaction trajectories were not sufficiently linear for robust initial-rate estimation, we did not derive percent inhibition. The separation between the matched trajectories nevertheless showed that modification-dependent recognition enhanced the activity of the appended inhibitory fragment.

We also explored 3-nitrotyrosine (3-NT), an oxidative tyrosine modification, in the design of HER2 binders (*29*). No experimentally determined HER2 complex containing 3-NT was used to define the designed interface. Of 16 experimentally tested designs, two showed detectable binding. Binder 5971 bound HER2 with a fitted K_D_ of 236.0 ± 4.3 nM, whereas the matched Tyr variant bound 3.3-fold more weakly (K_D_ = 780.4 ± 25.6 nM). Binder 5590 showed the same trend, with K_D_ values of 611.1 ± 12.6 nM and 1550 ± 47 nM for the 3-NT and Tyr variants, respectively (fig. S4). These matched substitutions show that the noncanonical nitrophenol group contributed favorably to recognition in interfaces generated without a pre-existing HER2–3-NT structural template.

### Noncanonical amino acids encode switchable and covalent binding

We next extended the framework to synthetic ncAA chemistries that enable post-production affinity control and covalent capture. These residues can also be incorporated by genetic code expansion at a defined site (*10, 11, 30*). We used this route to program three distinct molecular behaviors (Fig. 4). First, a tetrazine-bearing phenylalanine (Tet) was placed within an IL-7Rα interface (Fig. 4A). Nine of twelve selected binders bound IL-7Rα (fig. S15). A representative Tet-binder bound with a K_D_ of 8.4 ± 0.1 nM, whereas replacing Tet30 with Ala or Phe reduced affinity beyond 2 μM (Fig. 4, B and C). Thus, the noncanonical side chain contributed more than 200-fold to binding and acted as a chemically addressable determinant of the interface.

**Fig. 4.**
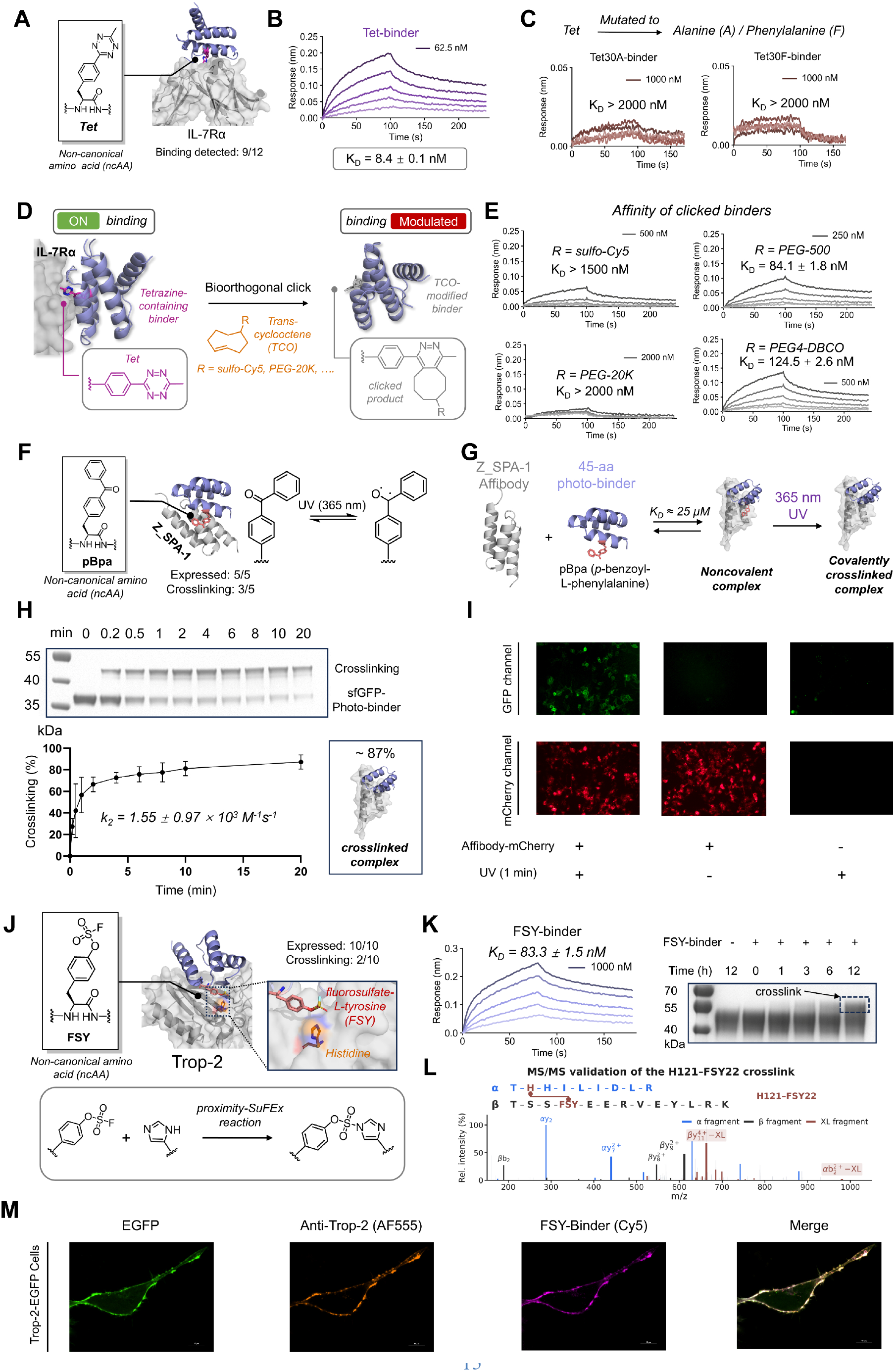
CheMoDesign programs ncAA chemistries into tunable and covalent protein binders. (A to C) A Tet-containing IL-7Rα binder. Nine of 12 designs bound IL-7Rα. The representative binder had KD = 8.4 nM; Tet30A and Tet30F had KD >2 μM. (D and E) TCO ligation installed distinct interface payloads and produced payload-dependent affinity changes. (F to I) A 45-residue pBpa photo-binder against Z_SPA-1. All selected designs expressed and three crosslinked. Irradiation at 365 nm produced approximately 87% crosslinking after 20 min (n = 3 independently performed experiments). Representative cell images were acquired after incubation with 10 μM photo-binder–sfGFP, 1 min of irradiation, and three DMEM washes. (J to M) FSY was oriented toward Trop-2 His121. All 10 designs expressed and two crosslinked. The selected binder had KD = 83.3 nM and accumulated crosslinked product from 0 to 12 hours; cross-linking MS supported the FSY22–His121 linkage. Confocal images show Cy5-binder colocalization with Trop-2 (scale bar, 10 μm). BLI uncertainties are fitting errors.

Tetrazines react rapidly and selectively with strained trans-cyclooctenes through the bioorthogonal inverse-electron-demand Diels-Alder reaction (*31, 32*). Tetrazine-TCO ligation therefore allowed the installed side chain to be rewritten after protein production (Fig. 4D). Reaction with TCO derivatives bearing sulfo-Cy5, PEG-500, PEG-20K, or PEG4-DBCO (fig. S5) produced distinct affinity states: K_D_ >1500 nM, 84.1 ± 1.8 nM, >2000 nM, and 124.5 ± 2.6 nM, respectively (Fig. 4E). The interface thus translated the identity and steric reach of a clicked payload into a change in target binding, including near-complete blocking by PEG-20K.

Second, we designed a 45-residue binder containing p-benzoyl-L-phenylalanine (pBpa) against the Z_SPA-1 affibody (Fig. 4F). Upon 365-nm irradiation, the benzophenone group of pBpa enters a reactive excited state that can covalently capture proximal C-H bonds (*33*). Five of five selected designs were expressed and three crosslinked the target (fig. S16). The selected design formed a weak noncovalent complex (fig. S6) but underwent rapid light-dependent crosslinking, reaching approximately 87% product after 20 min (Fig. 4, G and H, and fig. S7). On 293T cells displaying Z_SPA-1-mCherry, incubation with 10 μM photo-binder–sfGFP followed by 1 min of irradiation and three washes with DMEM produced target-dependent sfGFP retention (Fig. 4I). These representative images show that a low-affinity encounter complex can be converted into stable capture by a designed photochemical reaction.

Third, we oriented fluorosulfate-L-tyrosine (FSY) toward His121 of Trop-2 using a staged covalent-waking workflow (fig. S8). Aryl fluorosulfates can undergo proximity-enabled sulfur(VI) fluoride exchange with appropriately positioned nucleophiles and have been used to create covalent protein interactions (*34, 35*). All 10 designs were expressed, and two crosslinked Trop-2 (fig. S17). The selected FSY-binder bound noncovalently with a K_D_ of 83.3 ± 1.5 nM and accumulated covalent product over 0 to 12 hours (Fig. 4, J and K). Cross-linking mass spectrometry supported the designed FSY22–His121 linkage (Fig. 4L), and Cy5-labeled FSY-binder colocalized with Trop-2 on 293T cells (Fig. 4M). Together, the pBpa and FSY examples show that CheMoDesign can position mechanistically distinct ncAAs for direct photochemical capture or spontaneous proximity-enabled covalency.

### Post-expression chemistry couples protein recognition to new mechanisms

Genetic code expansion still limits designs to chemical groups that can be synthesized, transported, and incorporated by a compatible translation system. Post-expression conjugation, already widely used to manufacture chemically modified proteins, can install a broader range of structures and functions that are inaccessible to ribosomal encoding (*36*). It therefore brings the design process closer to the user-defined chemical-component principle of CheMoDesign: a canonical protein is generated around an arbitrary chemical group and then completed synthetically (fig. S9). We first targeted carbonic anhydrase IX (CA IX), a zinc metalloenzyme. Benzenesulfonamide provides a zinc-coordinating geometry suited to the CA active-site environment (*37*); a benzyl bromide reagent was therefore used to install a benzenesulfonamide-containing chemical modification (CMB) on a designed Cys (Fig. 5A). All 10 tested designs expressed and seven bound CA IX (fig. S18). The selected CMB-binder bound with a K_D_ of 45.4 ± 0.5 nM, whereas the unmodified precursor showed weak binding with a K_D_ of approximately 2.8 μM (Fig. 5B and fig. S10). Strong recognition therefore emerged from the designed combination of the protein and chemical components.

**Fig. 5.**
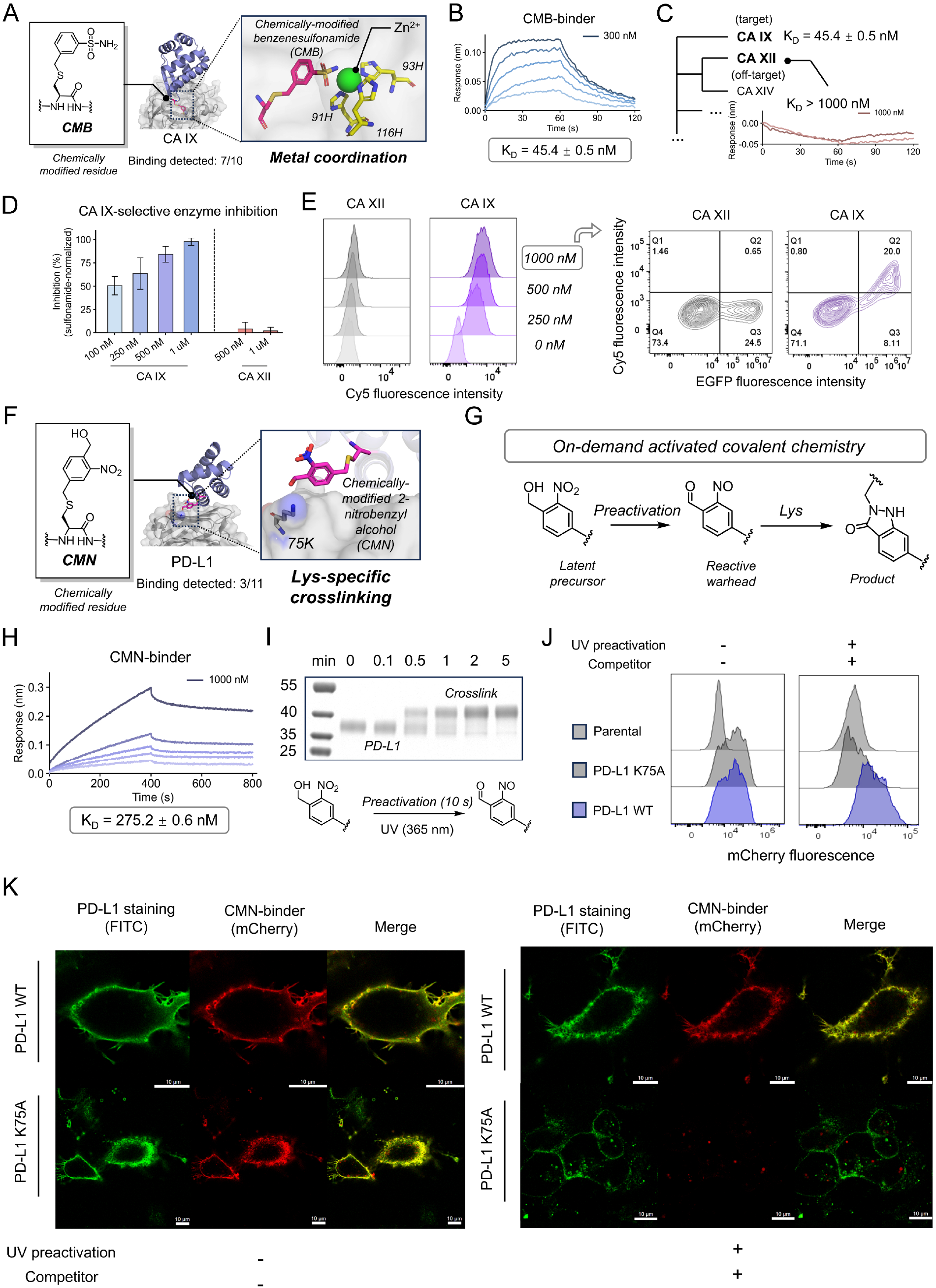
CheMoDesign extends protein design to post-expression chemical modification. (A to E) A benzenesulfonamide-containing modification (CMB) was installed on a designed Cys to target CA IX. Seven of 10 designs bound CA IX. The representative binder had KD = 45.4 nM, showed weak binding to CA XII (KD >1 μM), selectively inhibited CA IX in a pNPA esterase surrogate assay (mean ± SD, n = 3 parallel measurements), and bound CA IX-expressing cells. At 1 μM binder, inhibition was 97.8 ± 3.8% for CA IX and 2.2 ± 3.8% for CA XII. (F to K) A 2-nitrobenzyl alcohol-derived modification (CMN) was positioned toward PD-L1 Lys75. Three of 11 designs bound PD-L1 before modification. The representative binder had KD = 275.2 nM. Ten seconds of 365-nm preactivation generated an amine-reactive species that subsequently crosslinked PD-L1 (n = 3 independently performed experiments for gel-based crosslinking quantification). Competitor-resistant cellular retention occurred for PD-L1 WT but not K75A. Cell panels are representative of one experiment; confocal scale bars, 10 μm. BLI uncertainties are fitting errors.

The CMB group was modeled to enter the CA IX active site and coordinate catalytic Zn2+, while the surrounding generated protein interface contacted the isoform-specific surface (Fig. 5A). Consistent with this division of labor, the binder showed little detectable binding to the closely related CA XII at concentrations up to 1 μM (Fig. 5C) and selectively inhibited CA IX in a p-nitrophenyl acetate (pNPA) esterase surrogate assay (Fig. 5D). At 1 μM binder, CA IX activity was inhibited by 97.8 ± 3.8%, whereas CA XII inhibition remained near baseline at 2.2 ± 3.8% (mean ± SD, n = 3 parallel measurements). This near-complete, isoform-selective inhibition was obtained in a single design round without affinity maturation or experimental optimization. Flow cytometry likewise showed concentration-dependent binding to cells expressing CA IX-EGFP but minimal binding to CA XII-EGFP cells (Fig. 5E and fig. S11). Thus, the installed chemical group supplied catalytic-site engagement, while the generated protein surface conferred isoform discrimination.

Constitutively reactive electrophiles can impose challenges in chemoselectivity and reagent handling, motivating latent covalent warheads that are activated on demand (*38, 39*). We therefore sought an on-demand covalent binder that would remain latent during preparation and handling but acquire high crosslinking activity after a brief trigger. A 2-nitrobenzyl alcohol-derived modification (CMN), previously shown to generate an amine-reactive species upon photolysis (*40*), was installed on a designed Cys and positioned toward Lys75 of PD-L1 (Fig. 5, F and G). Three of 11 designs bound PD-L1 before modification (fig. S19). The selected CMN-binder bound with a K_D_ of 275.2 ± 0.6 nM (Fig. 5H). A 10-s pulse of 365-nm light delivered before target addition activated the latent warhead, and the preactivated binder rapidly formed a covalent complex when mixed with PD-L1 in PBS at 0.1 mg ml−1 of each protein, with t_1/2_ <1 min (Fig. 5I and fig. S12). Thus, activation was temporally separated from target encounter without sacrificing rapid capture.

We next tested whether this site-specific reaction could stabilize binding on cells. CMN-binder-mCherry (5 μM) was preactivated for 10 s, incubated with cells for approximately 10 min at room temperature, and then challenged for 40 min with 10 μM of the picomolar de novo PD-L1 binder deposited as PDB 8ZNL, which occupies the same epitope (*41*). After two DMEM washes, preactivated CMN-binder remained associated with wild-type PD-L1 despite competition, whereas retention was lost for PD-L1 K75A (Fig. 5J and fig. S13). Under the same conditions, confocal microscopy showed initial colocalization with both wild-type and K75A PD-L1, but competitor-resistant retention only with the wild-type protein (Fig. 5K). The K75A substitution therefore preserved detectable noncovalent recognition while removing the residue required for stable capture, supporting the designed Lys75 reaction. A brief light input could consequently be converted into durable, site-dependent labeling of a cell-surface antigen.

### Discussion and conclusion

CheMoDesign expands generative protein design from sequence and structure into chemical mechanism. Across the examples tested here, the noncanonical group was not merely tolerated by a pre-existing scaffold. Its removal, replacement, reaction, or activation altered the intended behavior: TYS strengthened thrombin recognition, Tet dominated IL-7Rα affinity and enabled post-design switching, pBpa and FSY produced orthogonal modes of covalent capture, CMB coupled metal coordination to CA IX selectivity and inhibition, and CMN converted light activation into delayed covalent reactivity. The common result is causal: the generated protein interface realizes a function that depends on chemistry specified before the interface is created.

The framework also suggests a route for transferring molecular recognition principles between domains that are usually designed separately. Protein–small-molecule complexes encode how synthetic functional groups interact with protein environments, whereas de novo protein design provides control over larger recognition surfaces. Combining these sources of information allows a compact chemical group to supply a mechanism and a generated protein to supply geometry, affinity, and selectivity. The CA IX binder illustrates this complementarity particularly clearly, although direct structural determination will be required to verify the modeled coordination and isoform-recognition geometry.

These results provide a foundation for extending chemically guided protein design. Experimental validation currently spans a defined set of targets and chemistries, with success rates varying across campaigns. The chemical space accessible in practice also depends on available atom-level representations and compatible routes for ncAA incorporation or post-expression modification. Although most designed complexes await atomic-resolution structures, mutational, biochemical, and cross-linking evidence supports the contribution of the designed chemistry; the precise contacts inferred from computational models remain to be verified. Expanding the chemical vocabulary, improving prospective calibration, and coupling design to higher-throughput experimental feedback should increase reliability and broaden the range of accessible mechanisms.

Natural evolution searches for function using chemistry that biosynthesis makes available. CheMoDesign inverts that order: a designer chooses a chemical capability, and the model generates a protein interface around it. This shift makes the chemical alphabet itself programmable and opens a design space in which molecular recognition, reactivity, and external control can be specified together. Because the framework operates entirely through a pretrained all-atom structure-prediction model, without task-specific training or fine-tuning, improvements in these models—particularly in protein-small-molecule recognition—could enable broader and more reliable design with noncanonical chemistry. We anticipate that this coupling will accelerate the development of protein reagents and therapeutics whose mechanisms cannot be reached within the canonical amino acid alphabet alone.

## Supporting information

supporting informations

## Acknowledgments

This work was supported by the National Natural Science Foundation of China (22325701 and 22637002 to T.L.), the Beijing Natural Science Foundation (Z250009 to T.L.), the Lingang Laboratory Grant (LGL-2615-01 to T.L.) and the NSFC Youth Student Basic Research Project (Doctoral Student) (224B2702 to Y.L.). We thank the High-Performance Computing Platform of Peking University Health Science Center for computational support.

## Data and code availability

All data supporting the findings of this study, except the exact construct sequences, are included in the manuscript and Supplementary Materials. The exact construct sequences and the full CheMoDesign source code will be released publicly soon (https://github.com/Yuxuan-Lee/CheMoDesign). Third-party software, model weights, and associated resources are subject to their original licenses and are not redistributed.

## Competing interests

The authors declare no competing interests.

## Notes

### Competing Interest Statement

The authors have declared no competing interest.

