## supporting informations for "De novo protein design in an expanded chemical space"

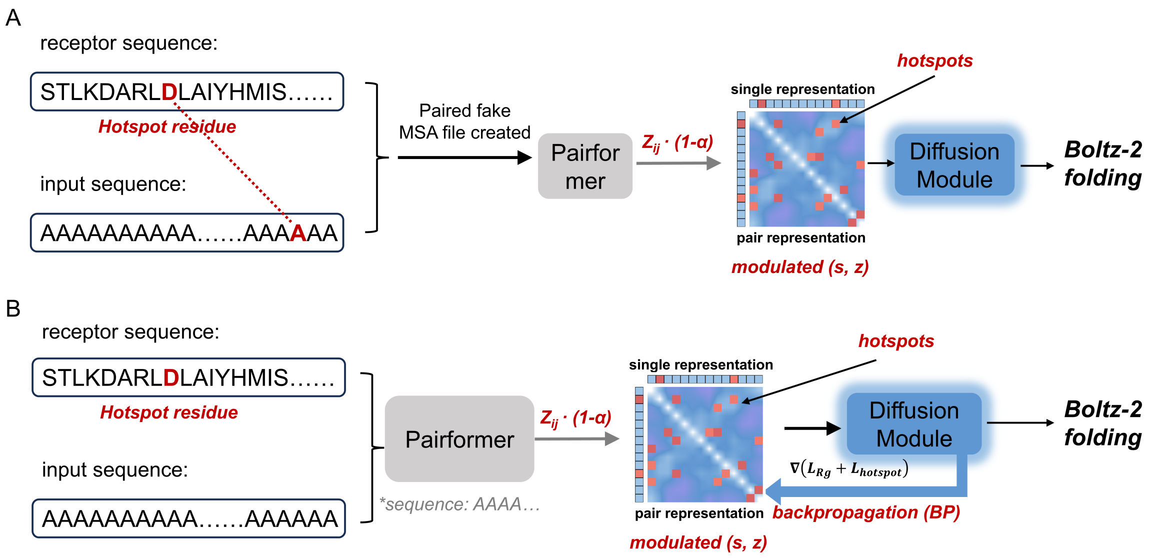


**Fig. S1. Two hotspot-guided strategies for retaining local interface constraints under relaxed global conditioning.**

(A) Schematic of hotspot guidance through artificial MSA-derived pair features ("fake MSA"). Using IL-7Rα as an example, designated receptor hotspot residues were treated as functional anchors. To introduce hotspot guidance, an artificial paired MSA was constructed by randomly selecting a hotspot residue from receptor and a binder residue pair, thereby introducing an additional bias into the Pairformer-derived pair representation. During representation scaling, pair features involving the hotspot residue were exempted from global attenuation $z_{ij} \rightarrow (1 - \alpha)z_{ij}$, whereas the remaining pair features were scaled. The resulting modulated single and pair representations *(s, z)* were then passed to the diffusion module for Boltz-2 folding. (B) Schematic of hotspot guidance through geometric-loss backpropagation (BP) without fake MSA. As in (A), pair features associated with the hotspot residue were preserved during scaling as functional anchors. Additional hotspot guidance was imposed by backpropagating a geometric loss composed of compactness and hotspot terms during diffusion, thereby biasing generated backbones toward the specified local interface geometry before Boltz-2 folding. Together, these strategies were used to preserve local interface requirements while relaxing global structural conditioning.

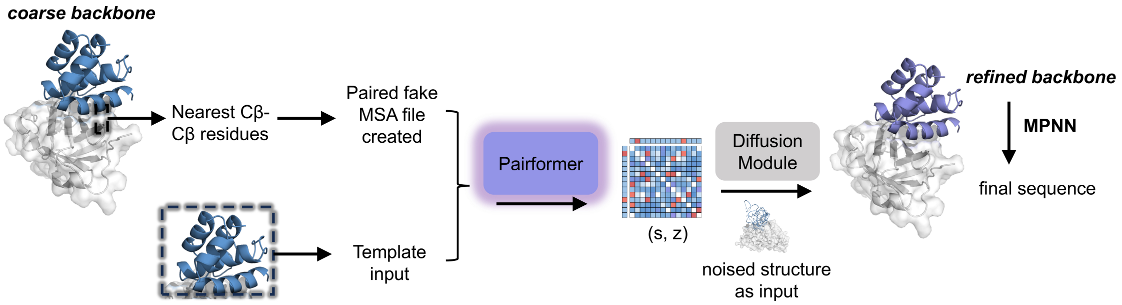


**Fig. S2. Schematic illustration of the sequence-conditioned waking step.**

Following the dreaming stage and initial MPNN sequence design, the workflow entered a waking step to refine local geometry. A coarse dreamed backbone was first used to identify nearest receptor–binder Cβ–Cβ residue pairs, which were then used to construct a paired fake MSA. In parallel, the coarse binder backbone was provided as a template input. Together with the designed sequence and a partially noised version of the dreamed backbone, these features were supplied to the Pairformer and a truncated diffusion denoising module. Unlike the hotspot-guided strategy described in Fig. S1, no explicit scaling or manipulation of $z_{ij}$ was applied in this waking step; guidance was instead provided through the binder-directed fake MSA, template features, and the truncated diffusion schedule. The waking step generated a refined backbone, which was subsequently subjected to final sequence design by MPNN.

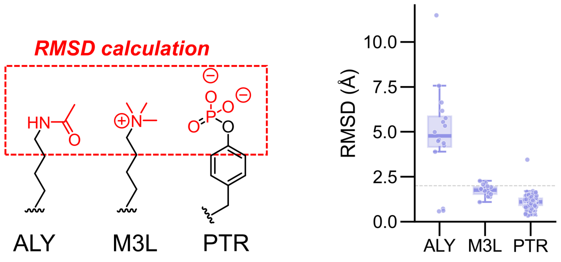


**Fig. S3. Recovery of native-like chemical-group geometry in PTM-containing designs.**

Left, chemical structures of ALY, M3L, and PTR, with the atoms used for functional-group RMSD calculation highlighted in red. Right, distributions of functional-group RMSD values for computationally passing designs generated for each PTM-containing complex. Each point represents an individual passing design. Using a functional-group RMSD threshold of 2 Å, native-like chemical geometry was recovered in 3 of 16 ALY designs, 14 of 17 M3L designs, and 122 of 123 PTR designs.

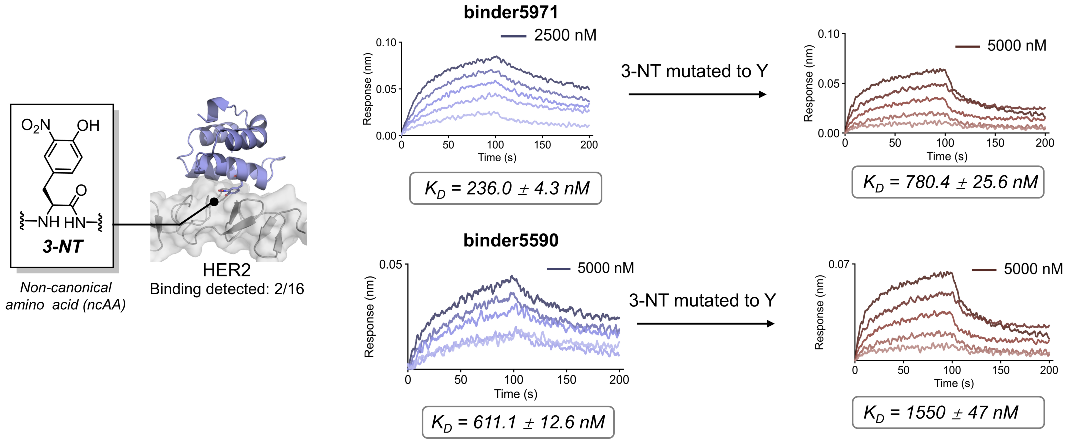


**Fig. S4. Binding characterization of 3-nitrotyrosine-containing HER2 binders.**

BLI characterization of designed HER2 binders containing 3-nitrotyrosine (3-NT) at the indicated interface position. Of 16 experimentally tested designs, two showed detectable HER2 binding. Representative sensorgrams are shown for binders 5971 and 5590 together with their corresponding Tyr-substituted variants, in which 3-NT was mutated back to canonical Tyr. Binder 5971 exhibited a fitted $K_D$ of 236.0 ± 4.3 nM with 3-NT, compared with 780.4 ± 25.6 nM after substitution with Tyr. Binder 5590 showed $K_D$ values of 611.1 ± 12.6 nM and 1550 ± 47 nM for the 3-NT-containing and Tyr-substituted variants, respectively. These measurements show that the designed binders retained HER2 recognition and that 3-NT contributed favorably to binding in both examples.

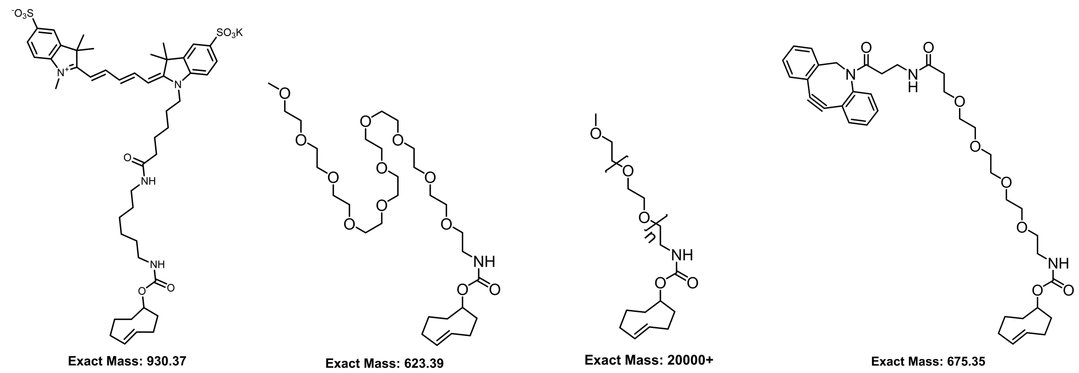


*TCO-sulfoCy5* *TCO-PEG500* *TCO-PEG20K* *TCO-PEG4-DBCO*

**Fig. S5. TCO derivatives used for post-translational rewriting of tetrazine-containing binders.**

Chemical structures of the trans-cyclooctene (TCO) derivatives used for tetrazine–TCO ligation: TCO-sulfoCy5, TCO-PEG500, TCO-PEG20K, and TCO-PEG4-DBCO. The corresponding conjugates were used to install chemically distinct side chains onto tetrazine-containing binders after protein production.

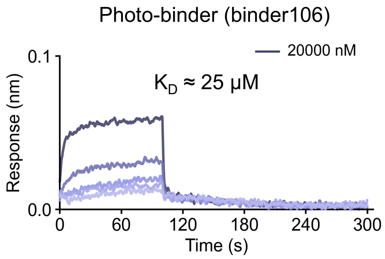


**Fig. S6. Noncovalent binding of the photo-crosslinking binder to its target.**

BLI characterization of photo-binder 106 before light-induced crosslinking. The binder showed weak noncovalent target binding, with an estimated $K_D$ of approximately 25 μM. This weak preassociation nevertheless supported efficient light-dependent covalent crosslinking described in Fig. 4.

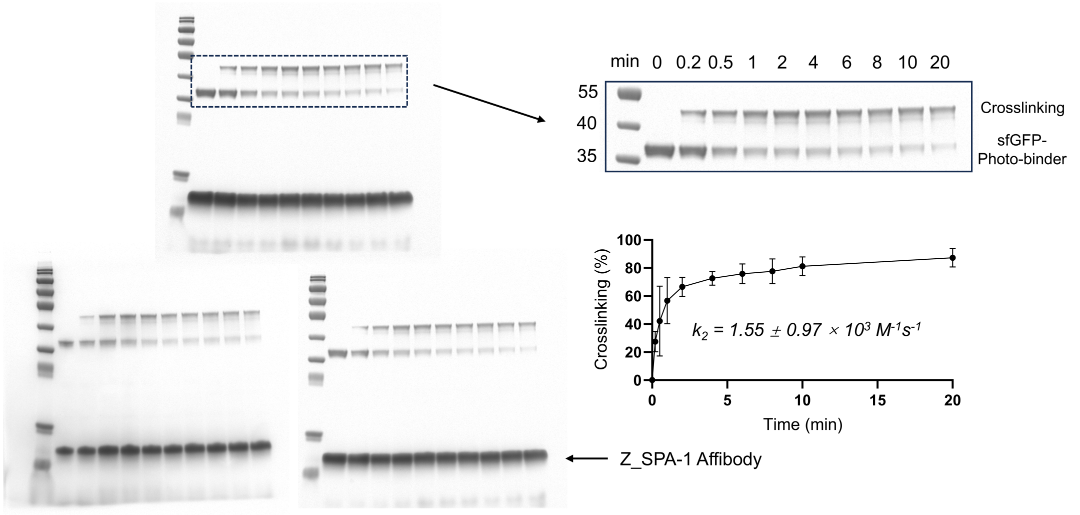


**Fig. S7. Full SDS-PAGE analysis and quantification of light-induced crosslinking by photo-binder 106.**

Full SDS-PAGE images from the time-course crosslinking experiment are shown, with the region containing the sfGFP–photo-binder and crosslinked product enlarged for clarity. Samples were collected after 0–20 min of light irradiation. The upper band corresponds to the crosslinked product, whereas the lower bands correspond to unreacted sfGFP–photo-binder and Z_SPA-1 affibody. Crosslinking efficiency was quantified from the indicated gel region and plotted as a function of irradiation time. Kinetic fitting gave an apparent second-order rate constant $k_2 = (1.55 \pm 0.97) \times 10^3$ $M^{-1}s^{-1}$.

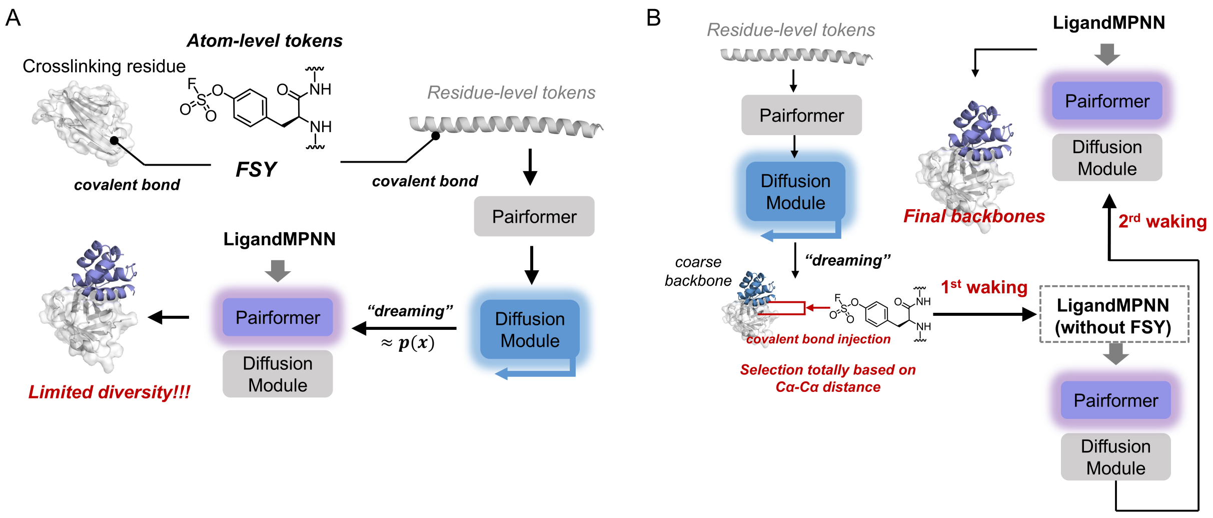


**Fig. S8. Staged covalent waking workflow for FSY-containing Trop-2 binders.**

(A) Initial strategy in which fluorosulfate-L-tyrosine (FSY) and its covalent linkage to the designated Trop-2 residue were explicitly encoded during backbone generation. Introducing the atom-level FSY constraint at a fixed binder position from the beginning strongly restricted backbone exploration and resulted in limited structural diversity. (B) Revised staged workflow that separates unconstrained backbone dreaming from covalent geometry refinement. A coarse binder backbone was first generated using residue-level representations without an explicit FSY crosslink. Candidate FSY positions were then selected solely according to a predefined Cα–Cα distance range between binder residues and the targeted Trop-2 His121 residue. FSY and the corresponding covalent linkage to His121 were subsequently introduced at the selected position. LigandMPNN was first used to design the surrounding binder sequence without explicitly including the FSY crosslink, followed by a covalent waking step in which the sequence-conditioned backbone was refined in the presence of the atom-level FSY linkage. Because introduction of the covalent constraint could shift the refined backbone relative to the initial sequence-conditioned structure, an additional LigandMPNN redesign and second waking step were performed to adapt the sequence to the covalently refined geometry. The resulting structures were retained as the final FSY-containing backbones.

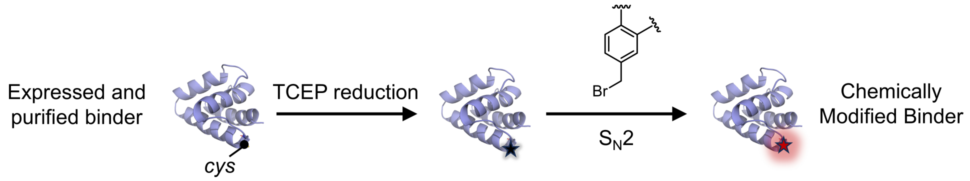


**Fig. S9. Post-expression chemical modification of designed binders through cysteine alkylation.**

Schematic of the chemical modification workflow used to install user-defined chemical groups onto designed binders after protein production. Purified binders containing a designated cysteine were first reduced with TCEP to generate a free thiol, followed by $S_{N2}$ alkylation with a benzyl bromide derivative bearing the desired chemical functionality. This reaction produced the final chemically modified binder at the predefined cysteine position.

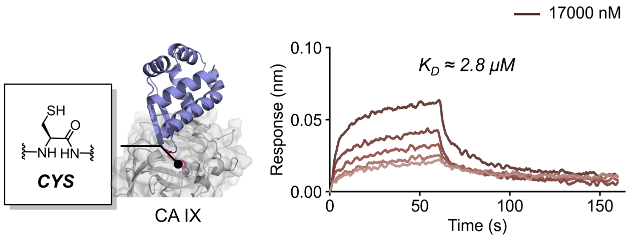


**Fig. S10. Binding of the unmodified Cys-containing precursor to CA IX.**

BLI characterization of the unmodified Cys-containing precursor of the CMB-binder. Before chemical modification, the precursor showed only weak binding to CA IX, with an estimated $K_D$ of approximately 2.8 μM. This weak baseline interaction contrasts with the substantially enhanced affinity of the chemically modified CMB-binder shown in Fig. 5B.

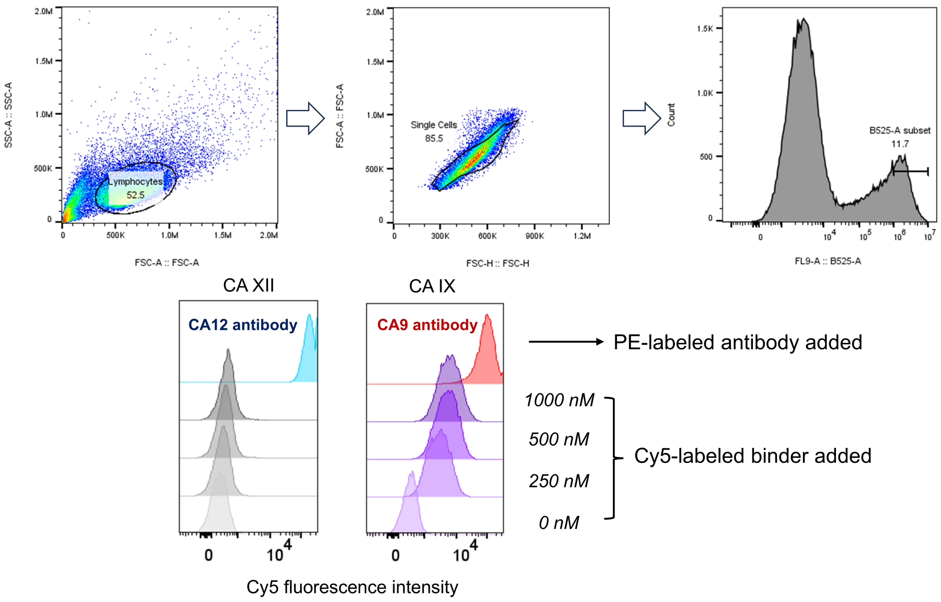


**Fig. S11. Flow-cytometry gating strategy and cellular binding of the CMB-binder to CA IX and CA XII.**

Top, representative gating strategy used for cell-based flow-cytometry experiments throughout this study. Cells were first gated by forward- and side-scatter characteristics, followed by singlet selection and subsequent gating of EGFP-positive cells to identify the population expressing the corresponding receptor. Bottom, binding of Cy5-labeled CMB-binder to cells expressing CA IX-EGFP or CA XII-EGFP at the indicated binder concentrations (0, 250, 500, and 1000 nM). PE-labeled antibodies against CA IX or CA XII were included as positive controls for receptor expression. The CMB-binder showed concentration-dependent binding to CA IX-expressing cells, whereas only minimal binding was observed for CA XII-expressing cells.

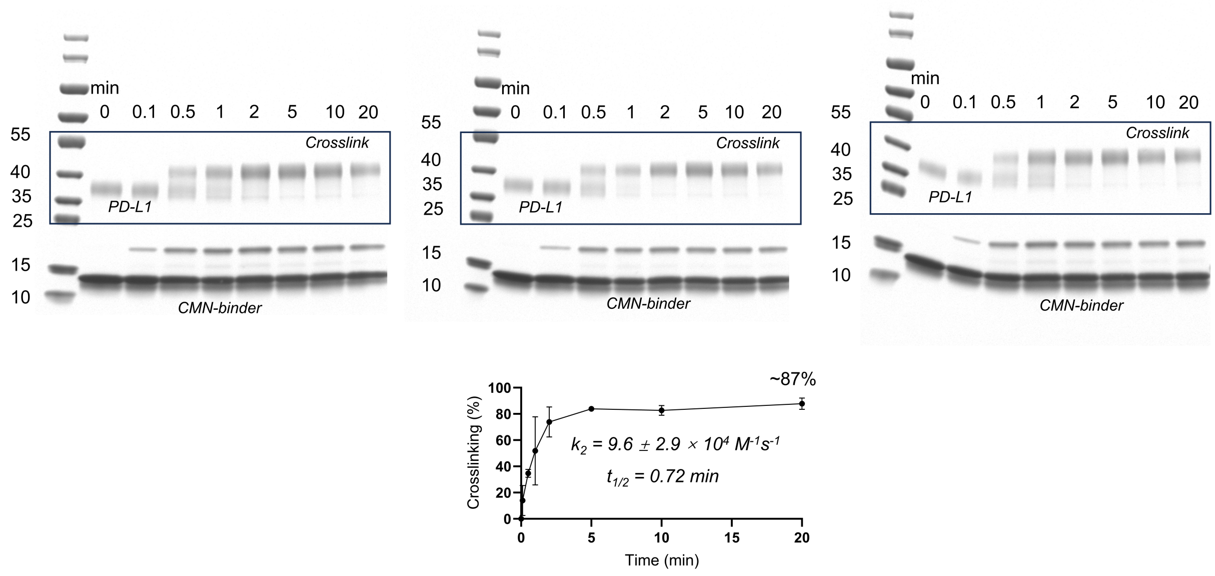


**Fig. S12. Full SDS-PAGE analysis and kinetics of covalent crosslinking between photoactivated CMN-binder and PD-L1.**

Full SDS-PAGE images from the time-course crosslinking experiments. CMN-binder was preactivated by a 10-s pulse of 365-nm light and subsequently mixed with PD-L1 in PBS at 0.1 mg/ml of each protein. Samples were collected over 0–20 min, with formation of the higher-molecular-weight covalent complex accompanied by depletion of the unreacted PD-L1 band. Quantification of the crosslinked product yielded an apparent second-order rate constant of $k_2 = (9.6 \pm 2.9) \times 10^4\ M^{-1}s^{-1}$, a half-time of $t_{1/2} = 0.72$ min, and approximately 87% crosslinking after 20 min.

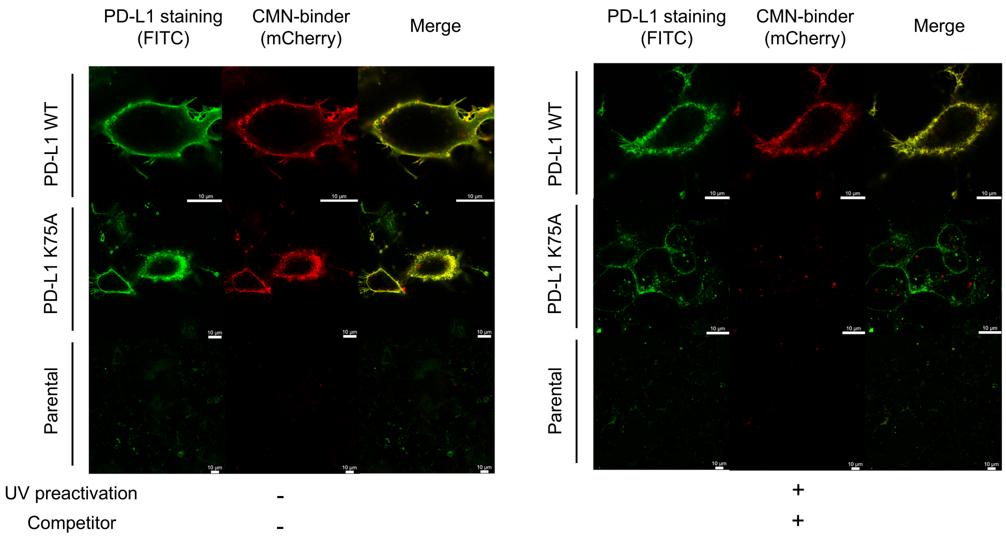


**Fig. S13. Confocal imaging controls for cell-surface labeling by CMN-binder.**

Representative confocal images of cells expressing wild-type PD-L1, PD-L1 K75A, or parental cells lacking exogenous PD-L1 expression. PD-L1 was visualized by FITC-antibody staining (green), and CMN-binder-mCherry is shown in red. Left, cells incubated with CMN-binder without UV preactivation or subsequent competitor challenge. Right, CMN-binder was preactivated by a 10-s pulse of 365-nm light before cell incubation, followed by challenge with the competing PD-L1 binder. CMN-binder colocalized with PD-L1 under noncompetitive conditions, whereas competitor-resistant retention after photoactivation was observed predominantly for wild-type PD-L1 and was strongly reduced for PD-L1 K75A. Parental cells showed minimal background signal under both conditions. Scale bars, 10 µm.

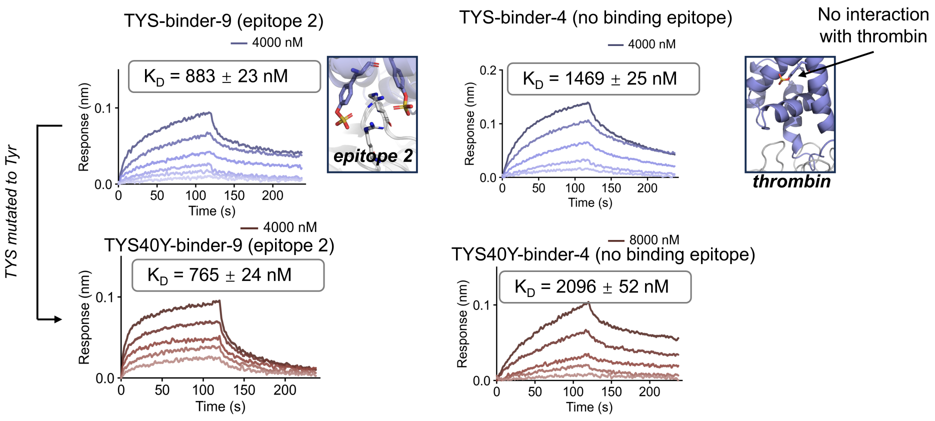


**Fig. S14. Binding characterization of additional sulfotyrosine-containing thrombin binders.**

BLI characterization of two additional thrombin binder designs containing sulfotyrosine (TYS) and their corresponding Tyr-substituted variants. TYS-binder-9 was designed against an alternative thrombin surface (epitope 2) and exhibited a $K_D$ of 883 ± 23 nM; substitution of TYS40 with Tyr resulted in a similar affinity $K_D$ = 765 ± 24 nM). TYS-binder-4 was designed without direct placement of TYS at the thrombin-binding interface and showed a $K_D$ of 1469 ± 25 nM. Substitution of TYS40 with Tyr reduced its affinity to 2096 ± 52 nM. Insets illustrate the designed binding geometry for TYS-binder-9 and the lack of direct interaction between the TYS-containing region of binder-4 and thrombin.

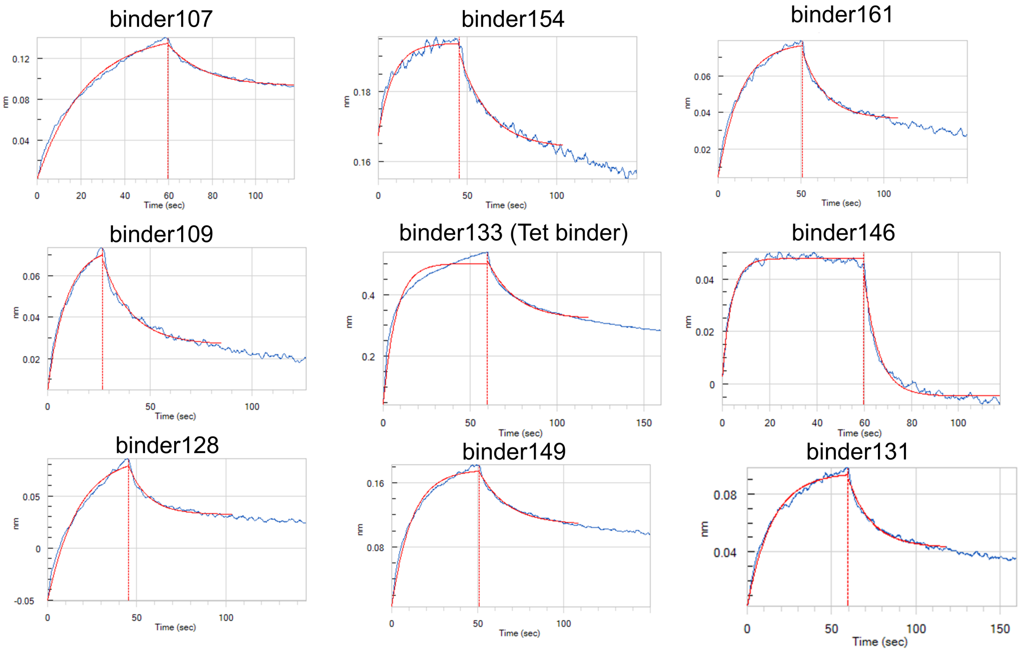


**Fig. S15. BLI screening of designed IL-7Rα binders.**

Representative BLI sensorgrams from the initial screening of designed IL-7Rα binders. Binding responses are shown for binders 107, 154, 161, 109, 133, 131, 146, 128 and 149. Binder 133 corresponds to the tetrazine-containing design selected for subsequent characterization. Blue traces show the experimental responses, and red curves show the fitted binding profiles.

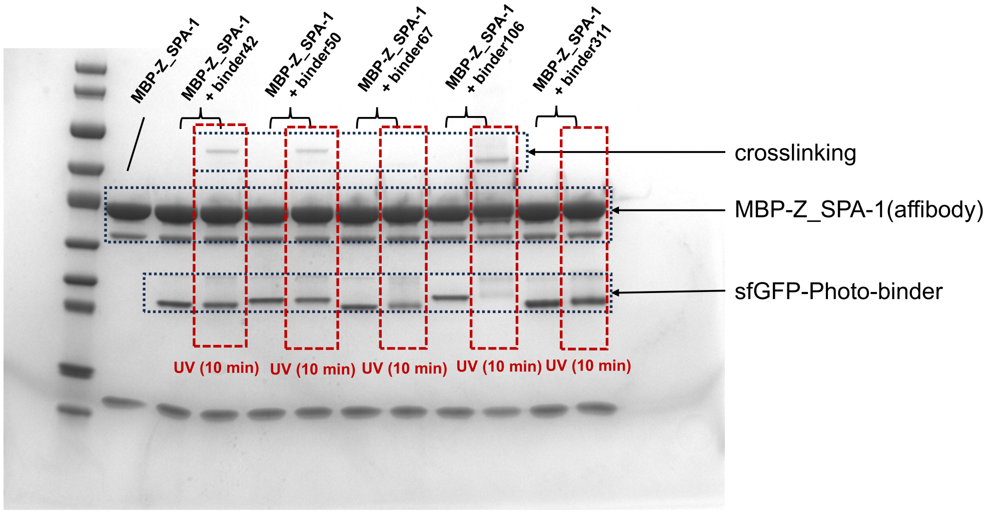


**Fig. S16. SDS-PAGE screening of photo-crosslinking pBPA-containing binders.**

SDS-PAGE analysis of UV-induced crosslinking for selected pBPA-containing binder designs fused to sfGFP and incubated with MBP–Z_SPA-1 affibody. Samples were analyzed with or without 10 min UV irradiation, and the positions of the MBP–Z_SPA-1 affibody, sfGFP–photo-binder, and higher-molecular-weight crosslinked product are indicated. Based on the appearance of the crosslinked band after UV treatment, three binder designs (binders 42, 50, and 106) were identified as positive candidates for target crosslinking.

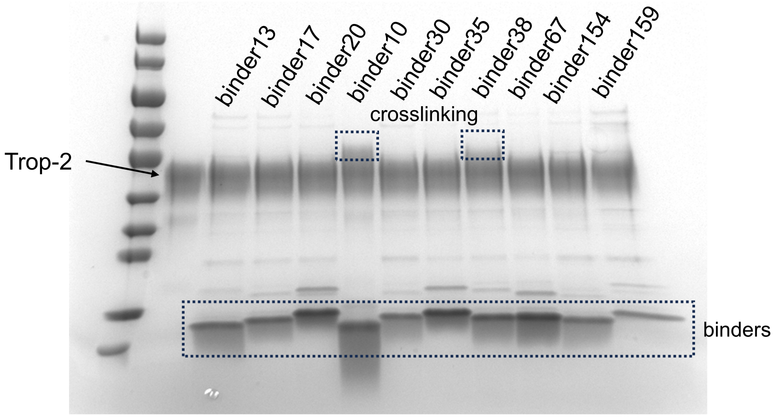


**Fig. S17. SDS-PAGE screening of FSY-containing Trop-2 binders for covalent crosslinking.**

SDS-PAGE analysis of designed FSY-containing binders incubated with Trop-2. The positions of free Trop-2, unreacted binders, and higher-molecular-weight crosslinked products are indicated. Among the tested designs, binders 10 and 38 showed detectable crosslinked products and were selected as positive candidates for subsequent characterization.

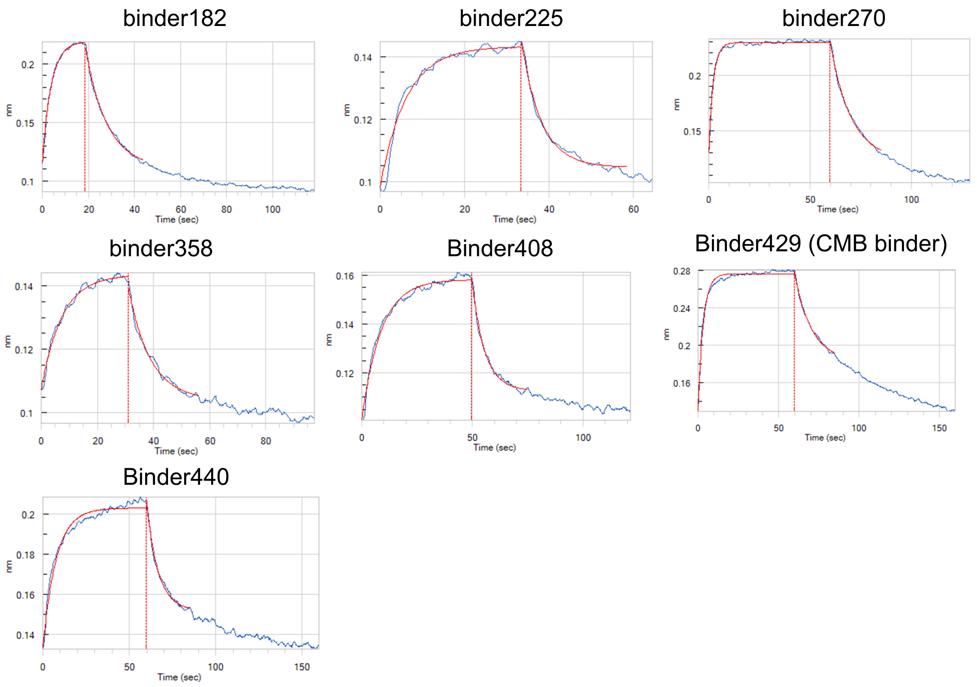


**Fig. S18. BLI screening of designed CA IX binders.**

Representative BLI sensorgrams from the initial screening of designed CA IX binders. Binding responses are shown for binders 182, 225, 270, 358, 408, 429, and 440. Binder 429 corresponds to the CMB-binder selected for subsequent chemical modification and functional characterization. Blue traces show the experimental binding responses, and red curves indicate the corresponding fitted binding profiles.

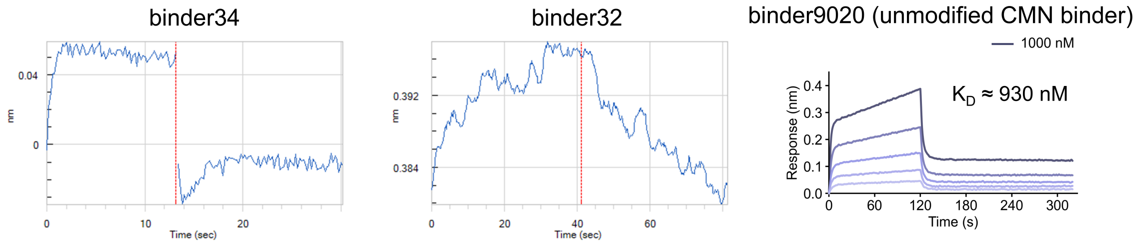


**Fig. S19. BLI screening of unmodified PD-L1 binder designs.**

BLI characterization of designed PD-L1 binders before chemical modification. Initial binding responses are shown for binders 34 and 32. Binder 9020, the unmodified precursor of the CMN-binder selected for subsequent chemical modification, was further characterized over a concentration series and exhibited an estimated $K_D$ of approximately 930 nM

### Materials and Methods

#### CheMoDesign implementation

CheMoDesign was implemented from the pretrained Boltz-2 all-atom structure-prediction model (v2.2.0; *22*). Model weights were frozen throughout. Pair representations were scaled as $z_{ij} \to (1-\alpha) z_{ij}$ with $\alpha = 0.9$ unless otherwise specified. Receptor–receptor entries and functional-anchor entries were exempted from scaling and were held fixed during gradient optimization. Internal single and pair representations were optimized by backpropagating a geometric loss (radius-of-gyration term, weight 1.0; helix term, weight 0.01; hotspot-contact term, weight 2.0) through one diffusion denoising step for 200 iterations using Adam *(42)* at a constant learning rate of 0.2 in FP32. Calculations used NVIDIA L40S GPUs (48 GB). Unless otherwise stated, Boltz-2 structure prediction used three recycling iterations. Target-specific diffusion schedules, scaling factors, anchor definitions, and loss settings are described below under target-specific configurations.

#### Functional-anchor construction

Functional anchors were defined from specified target hotspots or the atom-level chemical component of a PTM, ncAA, or post-expression modification. Residue-level and atom-level tokens were connected using the covalent-bond topology encoded by the corresponding CCD description *(43)*. All pair entries involving an atom-level chemical component, together with hotspot-associated pair entries, were exempted from scaling and were not updated during representation optimization. For hotspot-directed design, a passing backbone additionally required at least two binder residues within 7 Å by Cα-Cα distance of a specified target hotspot. Target-specific anchor definitions are described below under target-specific configurations.

#### Dreaming and waking

For the PD-L1 scaling experiment, 100 backbones were sampled at each α from an 80-residue polyalanine input. In the benchmarks preceding introduction of dreaming–waking (Fig. 2, A and B), four ProteinMPNN *(23)* sequences were generated per backbone for evaluation. For the five-target benchmark (Fig. 2D), each target generated 100 initial backbones (10 design rounds × 10 diffusion samples) from the same 80-residue affibody-derived starting sequence (AEAKYAKEMIFAVGEIYNLPNLTHGQIMAFIAALKDDPSQSSELLSEAKKLNDSQAPKAEAKYAKEAVGEIYNLGAAEFK). This repeated and truncated sequence was chosen to provide a protein-like composition and an α-helical prior relative to polyalanine. Before waking, LigandMPNN *(26)* (soluble_mpnn, sampling temperature T = 0.2) generated 10 sequences per coarse backbone, and the sequence with the highest overall_confidence score was retained. During waking, this sequence, a binder-directed synthetic paired MSA (fig. S2), template features from the dreamed backbone (forced template constraint, 3 Å), and dreamed coordinates at diffusion noise level $\sigma = 20$ were supplied to truncated Boltz-2 denoising. A 25-step schedule was restricted to steps with $\sigma \leq 20$, yielding approximately 10 denoising steps. Five woken structures were sampled per initial backbone. After a second backbone filter, LigandMPNN (soluble_mpnn, T = 0.2) generated 50 sequences per remaining refined backbone, and the two highest-confidence sequences were repredicted with AlphaFold 3 *(24)*.

The fake paired MSA is not an evolutionary alignment of the designed binder. Binders have no real homolog MSA; the construct exists only to give Boltz-2 a coevolution-like signal at the intended interface. Contact pairs were taken from the dreamed complex (Cβ/Cα within 8 Å). For each complex, a total of 4000 synthetic paired rows were generated and keyed with the same dummy taxonomy ID so Boltz-2 treats them as a paired MSA: with probability 0.8 the two contacting residues were mutated together under a complementary amino-acid map (charge–charge or hydrophobic–hydrophobic); otherwise they were mutated independently with BLOSUM62-like *(44)* conservative substitutions. Non-contact positions were mutated at 2%. The first row of each CSV is the query sequence itself. The fake receptor rows were concatenated with the receptor's real Jackhmmer *(45)* MSA; the binder MSA was entirely synthetic.

### Computational filtering and clustering

Backbones were filtered using loop content <40% and radius of gyration <14 Å. For RFdiffusion *(2)* designs against PD-L1, an additional hotspot-proximity filter (minimum two contacts within 7 Å of the designated hotspot residues) was omitted, because no RFdiffusion PD-L1 backbone passed it; CheMoDesign backbones and RFdiffusion designs against the remaining targets retained this filter. For the unrefined comparison branches of the five-target benchmark, candidate sequences were generated with LigandMPNN *(26)* (soluble_mpnn, T=0.2; 50 sequences per backbone, keeping the ten highest overall_confidence scores) without Rosetta relaxation. The post-waking branch retained two sequences per refined backbone, as described above. Selected candidates were repredicted with AlphaFold 3 *(24)*. For the computational benchmarks in Fig. 2, a predicted sequence–backbone pair was classified as passing only when minPAE ≤ 1.5 Å, ipTM ≥ 0.8, and binder pLDDT ≥ 80; minPAE was defined as the AlphaFold 3 chain-pair minimum PAE between the binder chain and the target chain (the chain_pair_pae_min summary entry for the binder–target pair; indices 0 and 1 in the input chain ordering). Every passing sequence–backbone pair was counted; sequences from the same backbone were not collapsed, and the number of unique originating backbones was reported separately. Structural clusters were assigned after computational filtering by complete-linkage hierarchical clustering at a TM-score cutoff of 0.6 using Foldseek *(46)* and TM-align *(47)* (exhaustive all-versus-all on the binder chain only; pairwise TM =min(qTMscore,tTMscore)). RFdiffusion designs were generated with RFdiffusion 1.1.0, checkpoint Complex_base_ckpt.pt, an 80-residue binder contig with target-specific hotspot residues, 50 diffusion steps, and default denoiser noise (noise_scale_ca = noise_scale_frame =1), and evaluated using the same sequence-design and AlphaFold 3 prediction pipeline as CheMoDesign.

### PTM recovery benchmark

Native complexes containing Nε-acetyllysine (ALY), Nε-trimethyllysine (M3L), or phosphotyrosine (PTR) were taken from reader domains of BRD4 bromodomain 2, CBX8 chromodomain, and SRC SH2, respectively (PDB 4KV4, 3I91, and 1SPS). Designs were initialized from an 80-residue affibody-derived sequence with a covalent PTM analog attached at Gly30 (CCD codes KACU, K3ME, and PTYU during backbone generation; ALY, M3L, and PTR during AF3 evaluation). Design was directed toward receptor hotspots W26 (BRD4), E36 and I41 (CBX8), and Y59 and K60 (SRC) in the numbering of the input constructs. For each complex, 100 backbones were generated (10 rounds × 10 samples; $\alpha = 0.9$). Sequences were designed with LigandMPNN (T = 0.2): 10 sequences were sampled on each passing coarse backbone and the top-scoring sequence was retained; after waking, 50 sequences were sampled on each refined backbone and the top two were evaluated by the standard computational filters. Recovery required a passing design and a functional-group RMSD <2 Å after superposition of the receptor. For chemical-ablation controls, the PTM at binder position 30 was converted to its canonical parent residue (ALY/M3L → Lys; PTR → Tyr) and the same design was repredicted; backbone generation and sequence design were not repeated.

#### General materials

Ni–NTA Agarose Resin 6FF was obtained from Sangon Biotech. Unless otherwise specified, chemical reagents were purchased from BLD Pharm and recombinant target proteins from Sino Biological. Genetic code expansion systems and ncAA incorporation conditions are summarized in Table S1.

#### Cell culture

HEK293T cells were obtained from Thermo Fisher Scientific and maintained in Dulbecco's modified Eagle medium (DMEM) supplemented with 10% fetal bovine serum (FBS) at 37 °C with 5% $CO_2$ in a humidified incubator.

#### Gene construction and protein expression

Genes encoding designed binders were synthesized or assembled in a pET-22b-derived expression vector. Construct assembly used high-fidelity PCR with 2× Phanta Max Master Mix (Dye Plus; Vazyme), assembly with the pEASY Pro Seamless Cloning and Assembly Kit (TransGen Biotech), transformation into E. coli DH5α (Sangon Biotech), and Sanger sequencing (GENEWIZ). Genetically encoded ncAA positions were specified by an amber (TAG) codon. Unless otherwise stated, binders carried a C-terminal sortase recognition motif followed by a hexahistidine tag; the pBpa photo-binder was expressed as a C-terminal GGGS–sfGFP fusion followed by the sortase motif and hexahistidine tag. Proteins were expressed in Escherichia coli BL21(DE3). Cultures in 2×YT medium were grown to $OD_{600}$ approximately 1.0, induced with 0.2 mM IPTG, and incubated for 16 h at 25 °C. For genetic code expansion, amber-containing constructs were coexpressed with the corresponding orthogonal aminoacyl-tRNA synthetase/tRNA system in the presence of the specified ncAA. The system, plasmid, ncAA concentration, and reagent source for each chemical functionality are listed in the supplementary table of ncAA incorporation conditions (Table S1). Cells were harvested by centrifugation, resuspended in PBS, and lysed by sonication. Lysates were clarified by centrifugation, and the soluble fraction was applied to Ni–NTA Agarose Resin 6FF (Sangon Biotech). The resin was washed with PBS containing 20 mM imidazole, and proteins were eluted with PBS containing 500 mM imidazole. Purified proteins were buffer-exchanged into PBST, quantified by BCA assay, and used for initial BLI screening. Selected binders were subsequently expressed at larger scale. For binders that were unstable during purification, 5% (w/v) trehalose was included and buffer exchange was performed by overnight dialysis with three buffer changes at 6-h intervals, followed by ultrafiltration concentration. Exact construct sequences are not included in this preprint and will be released soon.

#### Post-expression chemical modification

Cys-containing precursor binders were reduced with 500 μM TCEP and 1 mM DTT for 10 min and then buffer-exchanged by ultrafiltration into PBS containing 5% (w/v) trehalose, reducing residual TCEP to approximately 50 μM and DTT to <100 μM. CMB and CMN reagents were prepared as 100 mM stocks in DMSO and added directly to the protein solution at room temperature. CMB installation was performed with 1.7 mM reagent for 30 min, whereas CMN installation was performed with 500 μM reagent for 30 min. Protein concentration did not exceed one-fifth of the small-molecule reagent concentration. Reactions were quenched by addition of DTT to 5 mM, and products were dialyzed against PBS containing 5% (w/v) trehalose with three buffer changes. Intact-protein mass spectrometry indicated approximately 90% modification for CMB and approximately 100% modification for CMN. For Tet-binder remodeling, purified protein was incubated overnight with 0.1 mM of the indicated TCO derivative in PBS. Excess reagent was removed by ultrafiltration, except that the PEG-20K product was purified by size-exclusion chromatography on an ÄKTA system with a Superdex 75 Increase 10/300 GL column, using PBS as the mobile phase at 0.5 ml min−1. Elution was monitored at 280 and 260 nm, and fractions corresponding to the protein-containing peak were collected.

#### Biolayer interferometry

BLI measurements were performed on an Octet RED96 instrument (Pall ForteBio) using streptavidin (SA) or Protein A biosensors at 25 °C in PBST. Initial screens used CA IX-Fc, IL-7Rα-Fc, PD-L1-AviTag, and HER2-Fc. Kinetic characterization used biotinylated thrombin, CA IX-Fc, IL-7Rα-Fc, PD-L1, HER2-Fc, CA XII-AviTag, and Trop-2-AviTag. Human thrombin was obtained from Prolytix (cat. no. QQ1119-0.1MG); all other recombinant target proteins were obtained from Sino Biological. Thrombin was biotinylated in-house with 20 μM NHS-biotin before immobilization on SA biosensors. Fc-tagged or AviTag-biotinylated receptors were immobilized on Protein A or SA biosensors, respectively, to a loading response >1 nm, and the binder was used as analyte. For PD-L1 kinetic measurements, binders were C-terminally biotinylated by engineered sortase A (mgSrtA)-mediated transpeptidation *(50)*. Purified His-tagged binders (50–100 μM) were incubated with 10 μM mgSrtA and 5 mM biotin-PEG7-amine (BLD Pharm; CAS 1334172-76-7) in PBS at 25 °C overnight. His-tagged mgSrtA and cleaved C-terminal His-tag fragments were removed by subtractive Ni–NTA chromatography using Ni–NTA Agarose Resin 6FF (Sangon Biotech). Flow-through and wash fractions containing biotinylated binders were collected, and excess small-molecule substrate was removed by centrifugal ultrafiltration. Biotinylated binders were immobilized on SA biosensors, with PD-L1 used as analyte. In primary screening, 5–10 μM binder producing a response >0.02 nm was provisionally classified as binding. Each provisional hit was retested against an unloaded biosensor under the same conditions; designs producing >0.02 nm on the unloaded sensor were excluded as nonspecific binders. Kinetic measurements used five analyte concentrations in a twofold serial dilution, spanning Cmax to Cmax/16. The highest concentration and the association and dissociation intervals for each measurement are indicated in the corresponding sensorgram panels. Curves were reference-subtracted using the corresponding ligand-loaded biosensor exposed to PBST, aligned, and globally fit to a 1:1 binding model using ForteBio Data Analysis v11.0. Reported uncertainties are fitting errors unless otherwise stated.

#### Thrombin inhibition assay

Thrombin inhibition was evaluated using a chromogenic substrate assay. Human thrombin (Prolytix, cat. no. QQ1119-0.1MG) was diluted in assay buffer containing 50 mM Tris–HCl (pH 8.0) and 50 mM NaCl. Thrombin and chromogenic substrate were prepared as 4× stock solutions before use. Thrombin was preincubated with 5 µM TYS-binder-8-F7, the corresponding TYS40Y-binder-8-F7 variant, or 5 µM bivalirudin at 37 °C for 10 min. The final thrombin concentration was 1.4 nM. The reaction was initiated by addition of prewarmed N-(p-tosyl)-GPR-pNA acetate (TargetMol, cat. no. T36951) to a final concentration of 100 µM. Substrate hydrolysis was monitored at 405 nm every 20 s for 20 min using a microplate reader. Because substrate turnover was nonlinear over the measurement window, inhibition was compared using the complete OD405 kinetic profiles rather than a single inhibition value derived from fitted initial rates. Measurements were performed in three technical replicates, and data are presented as mean ± SD. No independent biological replicates were performed.

#### Photocrosslinking and proximity-enabled crosslinking

For pBpa photocrosslinking, 1 µM photo-binder–sfGFP and 10 µM Z_SPA-1 were mixed in PBS and irradiated at 365 nm using an 18-W UV lamp. Samples were held in microcentrifuge tubes on ice at approximately 1–5 cm from the lamp and irradiated for the indicated times. For the cell-surface pBpa assay, Z_SPA-1–mCherry-expressing 293T cells were incubated with 10 µM photo-binder–sfGFP in culture medium, irradiated for 1 min, washed three times with DMEM, and imaged directly. For FSY crosslinking, 10 µM FSY-binder and 2.5 µM Trop-2 were incubated in PBS at room temperature for 0 to 12 h. For CMN activation, 20 µl CMN-binder at 0.2 mg ml−1 in PBS was irradiated at 365 nm for 10 s on ice, then mixed with 20 µl PD-L1 at 0.2 mg ml−1 in PBS. The final reaction contained 0.1 mg ml−1 of each protein in a total volume of 40 µl. Reactions were sampled from 0 to 20 min and stopped by adding 10 µl protein loading buffer. Reaction products were resolved by denaturing SDS–PAGE, stained with Coomassie brilliant blue, and quantified by densitometry using ImageJ 1.49. For pBpa, the fraction crosslinked at each time point was calculated as 100% minus the intensity of the remaining free photo-binder–sfGFP band relative to the nonirradiated sample. For CMN, crosslinking was calculated from depletion of the free PD-L1 band relative to the initial sample. Quantifications were obtained from three independently performed experiments (n = 3).

#### Cross-linking mass spectrometry

Protein concentration was measured using a NanoDrop spectrophotometer before SDS–PAGE. The FSY-binder–Trop-2 crosslinked complex was isolated by SDS–PAGE, and 10 µg protein was subjected to tryptic digestion. Peptides were separated on an UltiMate 3000 nano-UPLC system (Thermo Scientific) using an Acclaim PepMap RSLC C18 column (75 µm inner diameter × 250 mm) with a 120-min gradient at 0.300 µl min−1. Mobile phase A was 0.1% formic acid in water, and mobile phase B was 0.1% formic acid in acetonitrile. The LC system was coupled directly to an Orbitrap Fusion Lumos mass spectrometer (Thermo Scientific), operated in data-dependent acquisition mode using Xcalibur 4.1.50. Full MS scans covered approximately m/z 350–1500 and were followed by higher-energy collisional dissociation (HCD) MS/MS scans. Crosslinked peptides were identified using pLink 3.0.17 *(51)* against a database comprising the Trop-2 construct (chain A) and the FSY-containing binder (chain B), with reverse decoy sequences generated by pLink. FSY was represented as His in the search database, and a custom His–His linker was configured with a crosslink mass shift of +88.9578 Da. This His encoding was used only for the database search; the experimental binder contained FSY at position 22. Carbamidomethylation of Cys was specified as a fixed modification, and oxidation of Met as a variable modification. Precursor and fragment mass tolerances were both 20 ppm. Crosslinked spectrum matches (CSMs) were filtered at a 5% false discovery rate (FDR). The search assigned 131 CSMs across multiple tryptic peptide forms to the binder FSY22–Trop-2 His152 pair, corresponding to His121 in the target numbering used for design and figures; additional assignments involved the adjacent His153 (His122 in design numbering). Representative scan 70534 matched TSS(FSY)EERVEYLRK–THHILIDLR.

#### Cell-surface binding and imaging

HEK293T cells were transfected with constructs encoding Z_SPA-1–mCherry, Trop-2–EGFP, CA IX–EGFP, CA XII–EGFP, PD-L1 WT, or PD-L1 K75A using PEI at a DNA:PEI ratio of 1:3, using 20 µg plasmid DNA per 15-cm dish. Medium was replaced 6 h after transfection. Cells were analyzed 48 h after transfection. For the pBpa cell-surface assay, Z_SPA-1–mCherry-expressing cells were incubated with 10 µM photo-binder–sfGFP, irradiated at 365 nm for 1 min, washed three times with DMEM, and imaged. For CMN competition experiments, CMN-binder–mCherry was prepared at 10 µM, preactivated at 365 nm for 10 s, and incubated with cells in PBS for approximately 10 min at room temperature at a final binder concentration of 5 µM. Cells were washed three times with DMEM containing 10% FBS and then challenged with 10 µM of a picomolar de novo PD-L1 binder recognizing the same epitope for 40 min at room temperature in DMEM containing 10% FBS. Cells were washed twice with DMEM before flow cytometry or confocal imaging. Conditions for Trop-2 and CA IX/CA XII cellular binding assays were 5 µM binder in DMEM for 60 min on ice, followed by two DMEM washes; cells were kept on ice until analysis. Flow cytometry was performed using a CytoFLEX flow cytometer (Beckman Coulter) and analyzed with FlowJo 10.8.1; the gating strategy is illustrated in Fig. S11. Confocal images were acquired using a STELLARIS 8 microscope (Leica Microsystems) and processed using LAS X Office 1.4.7.28982. Cell images are representative of one experiment unless otherwise stated; scale bars are 10 µm.

#### Carbonic anhydrase inhibition

CA IX or CA XII (approximately 30 nM) was incubated with CMB-binder at 100, 250, 500, or 1000 nM in PBS at 37 °C for 10 min. p-Nitrophenyl acetate was added to 1 mM, and absorbance at 400 nm was monitored for 30 min using a microplate reader. Net rates were calculated from OD400 slopes after subtraction of matched no-enzyme blanks. Percent inhibition was normalized to the sulfonamide-inhibitable window, with no inhibitor defined as 0% and 1 mM benzenesulfonamide as 100%; negative benzenesulfonamide-corrected rates were set to zero before normalization, and displayed values were limited to 0–100%. Data are mean ± SD from three parallel measurements.

#### Candidate selection for experimental testing

The three-threshold definition above was used for the Fig. 2 computational benchmarks. For prospective experimental campaigns, ipTM ≥0.8 and pLDDT ≥80 were used as mandatory confidence thresholds; minPAE ≤1.5 Å was applied when sufficient candidates met all three criteria but was not required in campaigns yielding too few candidates. Passing designs were then inspected manually before experimental testing. Designs were excluded when they contained clearly buried charged residues (Arg, Lys, Asp, or Glu) in a hydrophobic core, presented an interface comprising only approximately three residue–residue contacts, or placed the specified ncAA or chemical component outside or incompatibly oriented within the intended interface. Inspection was stopped after 10–16 candidates had been accumulated for a campaign and rarely extended beyond 100 computationally passing designs. Candidate selection was completed before experimental testing, and no subsequent round of experimental optimization was performed.

#### Statistical analysis

No statistical method was used to predetermine sample size. Error bars and n definitions are stated in the corresponding figure legends. BLI uncertainties are fitting errors from 1:1 kinetic fits. Cell images are representative and were not used for quantitative statistical inference. Exact tests and P values are reported only for datasets comprising independent experimental replicates.

**Table S1.**

**Genetic code expansion systems and ncAA incorporation conditions.**

Genetic code expansion conditions for the E. coli expression experiments. Post-expression CMB and CMN conjugation is described separately in Methods.

| ncAA and application | aaRS and suppressor tRNA | Expression plasmid and selection | ncAA conc. | Source and reference |
|---|---|---|---|---|
| TYS Thrombin binders | M. jannaschii TyrRS-derived sTyrRS / Mj tRNA^Tyr_CUA | pULTRA-sY (Addgene 82417); spectinomycin, 50 µg ml−1 | 10 mM | BLD Pharm; CAS 98930-06-4; *(48)* |
| Tet2-Me IL-7Rα binders | M. jannaschii TyrRS-derived Tet-v2.0RS / Mj tRNA^Tyr_CUA | pDule2-Tet2.0 (Addgene 85497); spectinomycin, 50 µg ml−1 | 1 mM | BLD Pharm; CAS 2363129-95-5; *(31)* |
| pBpa Z_SPA-1 photo-binders | M. jannaschii pBpaRS / Mj tRNA^Tyr_CUA | pEVOL-pBpF (Addgene 31190); chloramphenicol, 25 µg ml−1 | 1 mM | BLD Pharm; CAS 104504-45-2; *(33)* |
| FSY Trop-2 binders | M. barkeri PylRS-derived FSYRS / Mb tRNA^Pyl_CUA | pEVOL-FSYRS; chloramphenicol, 25 µg ml−1 | 1 mM | BLD Pharm; CAS 2227199-79-1; *(34)* |
| 3-NT HER2 binders | M. alvus PylRS NYF5 / Ma tRNA^Pyl_CUA | pAJE-Ma pylRS nitroY/haloY-F5 (Addgene 225684); spectinomycin, 50 µg ml−1 | 2 mM | BLD Pharm; CAS 621-44-3; *(49)* |

TYS, sulfotyrosine; Tet2-Me, hydrochloride salt of (S)-2-amino-3-[4-(6-methyl-1,2,4,5-tetrazin-3-yl)phenyl]propanoic acid; pBpa, p-benzoyl-L-phenylalanine; FSY, fluorosulfate-L-tyrosine; 3-NT, 3-nitrotyrosine.
